# Autoregulatory circuit regulating basolateral cargo export from the TGN: role of the orphan receptor GPRC5A in PKD signaling and cell polarity

**DOI:** 10.1101/2020.05.26.114710

**Authors:** Rosaria Di Martino, Anita Capalbo, Lucia Sticco, Alessandra Varavallo, Vidya Kunnathully, Valentina De Luca, Namrata Ravi Iyengar, Matteo Lo Monte, Petra Henklein, Jorge Cancino, Alberto Luini

**Affiliations:** Institute of Biochemistry and Cell Biology, National Research Council, Via Pietro Castellino 111, 80131 Naples, Italy; Telethon Institute for Genetics and Medicine (TIGEM), Pozzuoli, Italy; Institut fur Biochemie, Charite Universitätsmedizin, Berlin, Germany; Centro de Biología Celular y Biomedicina (CEBICEM), Facultad de Medicina y Ciencia, Universidad San Sebastián, Lota 2465, Santiago 7510157, Chile

**Keywords:** Apico-basal polarity, autoregulatory systems, basolateral cargo, GPRC5A, PLCβ3, PKD, signaling, TGN export

## Abstract

The membrane transport apparatus comprises a series of separate membrane bound compartments, or transport stations, that are responsible for the synthesis, processing, transport, sorting and delivery to their final cellular destinations of most transmembrane and soluble lumenal proteins. Over the last decades the membrane transport system has been shown to be extensively regulated both by environmental inputs and by internal homeostatic signalling systems, or control systems, that operate to maintain the homeostasis and optimal functionality of the main transport stations, such as the endoplasmic reticulum and the Golgi, in the face of internal and external perturbations. The trans-Golgi network (TGN) is a major transport and processing station and the main sorting compartment of the transport apparatus. However, the mechanisms that control cargo export and sorting at the TGN have so far remained elusive. Here we focus on the sorting of basolateral cargo proteins and show that these proteins bind to the TGN localized orphan receptor GPRC5A. The cargo-GPRC5A complex triggers the activation of a signaling pathway that involves the Gβγ subunits dependent activation of the phospholipase C beta 3 (PLCβ3), which inturn induces diacyl glycerol (DAG) production. DAG recruits and activates protein kinase D (PKD) and the phosphorylation of its substrates. This step results in the formation of basolateral carriers for delivery of these cargoes to the basolateral plasma membrane domain. We term this mechanism “ARTG” (**A**uto**R**egulation of **TG**N export). Remarkably, the impairment of ARTG pathway components, and in particular of GPRC5A, causes defects in the polarized organization of epithelial cells.

## Introduction

The biosynthetic membrane transport system is a series of membrane-bound compartments arranged in a linear sequence that is responsible for the synthesis, processing, transport, sorting and delivery to their final cellular destinations of most transmembrane and soluble lumenal proteins (henceforth called cargoes), and of most lipids. It includes the endoplasmic reticulum (ER), where the initial synthesis and processing of cargoes occurs, the ER-Golgi intermediate compartment (ERGIC), the Golgi complex, and the trans-Golgi network (TGN). The TGN is considered to be the main sorting station from where cargo proteins are transported to their final destinations, such as lysosomes, the plasma membrane (PM) or the extracellular milieu (De Matteis and Luini, 2008).

Throughout their journey, cargo proteins are processed by specialized reactions (folding, followed by post-translational modifications like glycosylation, and then sorting) in successive stations, each of which resuts in the generation of intermediate products, such as unfolded or partially modified proteins, which are further processed and transported across the successive secretory stations (Mellman and Warren, 2000).

Over the last several years, a growing body of evidence has led to the recognition that membrane transport is supervised by multiple regulatory mechanisms, among which we can recognize two different types. One class of regulatory devices acts in response to external signals, and integrates the transport system to a variety of complex cellular responses initiated by PM receptors, such as such growth, migration or differentiation responses (Farhan and Rabouille 2011; Farhan et al., 2017). A second type of mechanism, called internal homeostatic regulatory devices, operate to monitor the optimal working of the transport compartments and fuction to A) maintain their homeostasis and efficiency in the face of internal and external perturbations, and B) ensure that the compartments act in a coordinated fashion amongst themselves and with other functional modules in cells, such as metabolism or cytoskeletal motility (Luini et. al., 2014; Luini and Parashuraman, 2016)

A well characterized homeostatic device is the unfolded protein response (UPR), which regulates the levels of unfolded proteins in the ER through multiple signaling and transcriptional cascades (Walter and Ron, 2011). Other examples are the autoregulatory systems that operate in the ER (Subramanian et al., 2019; Centonze et al., 2019) and the Golgi (Pulvirenti et al., 2008; Giannotta et al., 2012; Cancino et al., 2014; Solis et al., 2017; Tapia et al., 2019) that ensure the correct trafficking through these stations. So far, the organization, molecular determinants and function of a potential autoregulatory circuit at the TGN has not been examined.

When cargo proteins reach the lumen of the TGN they are sorted and shipped to either the basolateral or apical PM domains, or to the lysosomes. However, in the absence of a homeostatic mechanism, they may accumulate in the TGN lumen, creating the risk of ectopic and functionally aberrant signaling (Mellman and Nelson, 2008). Moreover, the accumulation of cargo in the TGN might result in their mis-sorting. Indeed, it is known that in a sorting organelle if the preferred export route for a certain cargo, for example a basolateral cargo protein, is blocked, the cargo proteins will end up leaving the organelle by means of apical transport carriers, with potentially harmful functional consequences (Stoops and Caplan, 2014; Wilson, 2011). Based on these considerations, we predict that an autoregulatory device is expected to operate at the TGN to regulate cargo sorting and export.

To reveal the presence of such a device, we challenged the transport apparatus with an overload of basolateral cargo in the TGN and analyzed the cellular response. First, we used a targeted phosphoproteomics approach to detect the modulation of signaling pathways, using an assay designed to detect the activation of a large number of kinases and other regulatory molecules. Second, to complement these findings, we undertook an unbiased proteomics approach to determine the interacting partners of basoloteral cargo accumulated at the TGN by immuno-precipitation and mass spectrometry (see Materials and Methods).

We report here that basolateral cargo proteins, upon arriving at the TGN, activate an orphan TGN resident G-Protein Coupled Receptor GPRC5A that is in complex with heterotrimeric G-protein subunits Gαi3 and Gβγ. GPRC5A acts as a sensor of cargo at the TGN, and activates a network of signaling pathways, thus forming a central component of the autoregulatory device. Among the activated cascades, GPRC5A simulates a Gβγ-PLCβ3-PKD signaling pathway that has been characterized previously to regulate export from the TGN (Jamora et al., 1999; Liljedahl et al., 2001; Baron and Malhotra, 2002, Diaz Añel and Malhotra 2005). Activated Gβγ subunits recruit and activate the enzyme phospholipase C beta 3 (PLCβ3), which produces the diacyl glycerol (DAG) required for protein kinase D (PKD) recruitment and activation. PKD activation results in phosphorylation of many substrates and in the formation of basolateral cargo carriers that export cargo out of the Golgi, attenuating basolateral protein cargo load in the TGN lumen and enhancing the fidelity of sorting. These series of events result in a negative feedback action on the concentration of cargo in the TGN, thus reducing the risk of aberrant cargo accumulation and mis-sorting.

We call this circuit ARTG (**<u>A</u>**uto**<u>R</u>**egulation of **<u>TG</u>**N export). Importantly, the ARTG signaling pathway supports the maintenance of cell polarity, and may control the development of the epithelial-mesenchymal transition (EMT).

## Results

### A cargo overload in the TGN triggers signaling responses that regulate cargo export

To challenge the autoregulatory device expected to operate at the TGN with an appropriate perturbation, we set up conditions to generate a basolateral cargo protein overload in this compartment using different transport synchronization methods. Such methods were based on the use of retention using selective hooks (RUSH) (Boncompain et al., 2012) and FM (Rivera et al., 2000; Rollins et al., 2000) constructs of basolateral cargo proteins, as well as the temperature-sensitive variant of the vesicular stomatitis virus (ts045) G protein (for brevity, VSVG), an extensively characterized basolateral cargo protein. These proteins can be accumulated in controlled amounts in the ER by blocking their ER export (see Methods for experimental details), and can then be allowed, by removing the export block, to exit the ER and move as a synchronous ‘wave’ through the successive stations of the secretory pathway, including the TGN. A well-characterized variant of this protocol is based on an additional temperature block at 20°C for 2 hours after release from the ER. At 20°C, cargo proteins accumulate in the TGN (Matlin and Simons, 1983). Upon shifting the temperature again to 37°C (or 32°C in case of VSVG) they leave the TGN for the PM. We call these two protocols the “wave” and the “20°C accumulation”, respectively.

The export of VSVG from the TGN has been extensively characterized before (Polishchuk et al., 2003; Luini et al., 2007) and has been shown to take place via large pleiomorphic carriers that form and deliver cargo to the PM at rates roughly proportional to the load of cargo in the TGN indicating that the rate of carrier formation reflects the TGN cargo load (De Matteis and Luini, 2008; Polishchuk et al., 2009). To uncover the mechanism responsible for coupling the rate of export to the amount of cargo load, we sought to identify the expected signaling responses using a targeted phosphoproteomic approach based on a phospho-antibody microarray. We first generated a VSVG traffic “wave” using previously characterized conditions i.e. by accumulating this protein at 40°C in the ER for 3 hours and then releasing the block by shifting the temperature to 32°C (Mironov et al., 2003). Cycloheximide (CHX) was added from 30 min before releasing the block until the end of the experiment to exclude effects from endogenous cargoes. VSVG exited the ER synchronously and reached its peak concentration in the TGN between 20 and 25 minutes and then reached the PM between 30-40 minutes after the release of the ER block. The phospho-antibody microarray was used to assess the phosphorylation state of several hundred kinases and regulatory proteins at the time of accumulating VSVG in the TGN (i.e., 20 minutes at 32°C after the release of the ER block) in comparison to when cargo accumulation in the ER at 40°C (Figure 1A). The TGN cargo peak induced the hyper-phosphorylation or de-phosphorylation of approximately 90 proteins on specific residues (Figure 1B). Analysis of this data-set using bioinformatic tools and manual curation, revealed the activation of various signaling pathways (Figure 1C). The most enriched were the ERK/MEK signaling, Calcium/DAG/PKC/PKD signaling, JAK/STAT, and mTOR pathways. In addition, we also noted an elevated phosphorylation of eIF2α, a mechanism linked to the inhibition of protein synthesis (Pakos-Zebrucka 2016; Costa-Mattioli and Walter, 2020).

**Figure 1:**
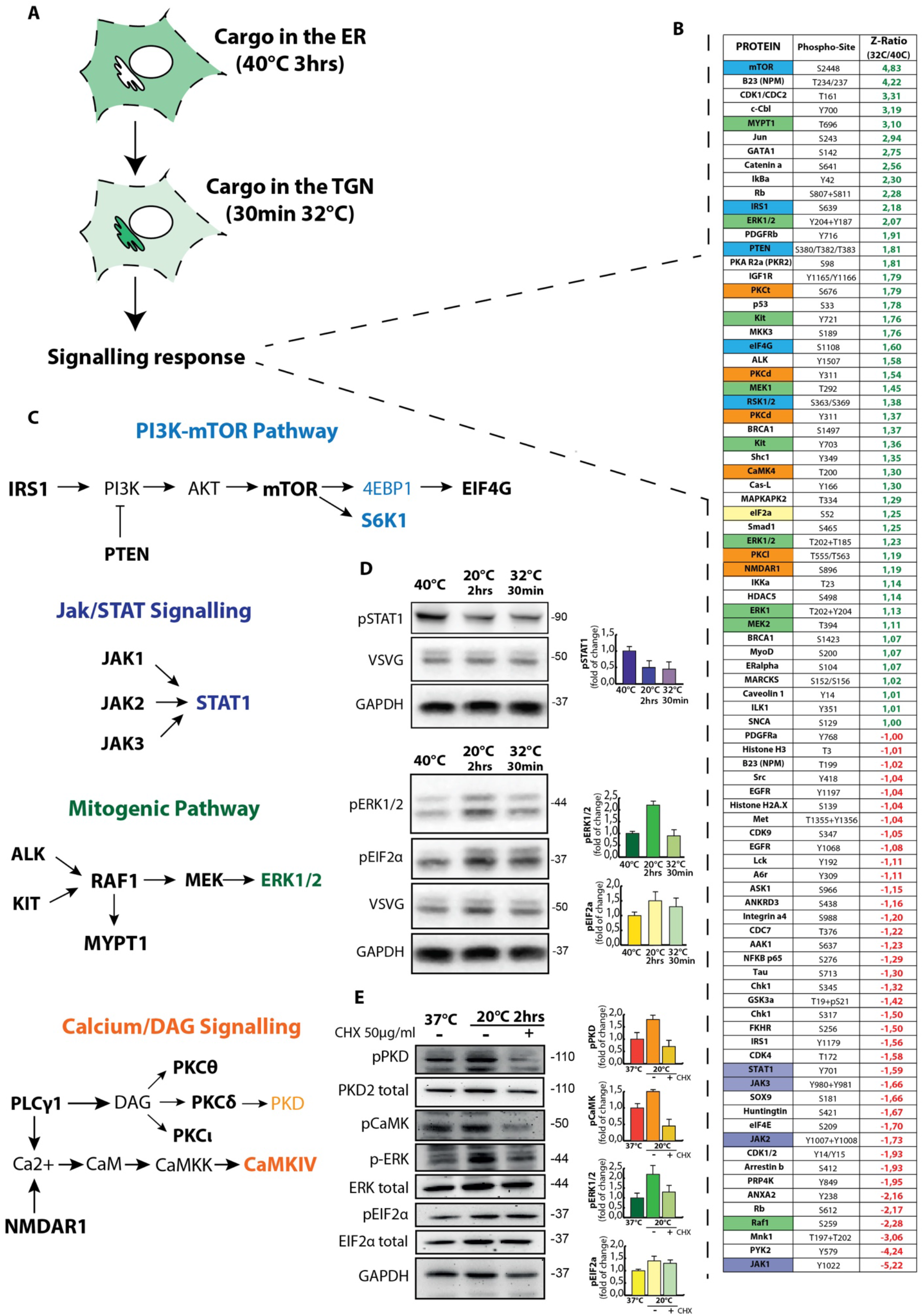
The phospho-antibody microarray reveals the signals activated by the arrival of cargo in the TGN. **A)** The global signaling response generated by the arrival of cargo (VSVG) in the TGN was analyzed through the phospho-antibody microarray made available by the company Kinexus. The two samples analyzed were the “Cargo in the ER” versus “Cargo in the TGN”. Briefly, after infecting the cells, they were collected at their respective timepoints and their cell pellets were sent to the Kinexus Company for analysis. **B)** Shortened list of results obtained from the analysis of the phospho-antibody array. The Z-ratio values greater than 1 and less than −1 were considered significant (in green the values that indicate an increase in phosphorylation and in red those that indicate a decrease). The cells in the protein name column are colored based on their belonging to specific signaling pathways: blue for the PI3K-mTOR pathway, green for the Mitogenic pathway, orange for the Calcium/DAG pathway and finally purple for the Jak-STAT signaling. **C)** The proteins obtained from the shortened list have been analyzed by Gene Ontology and manual curation in order to highlight particular signaling pathways activated or inactived by the arrival of the cargo in the TGN. This analysis highlighted the activation of the MAPK (Mitogenic), PI3K-MTOR and Calcium/DAG pathways, while the JaK-STAT pathway is inhibited. In the representation of the pathway, the following graphic codes have been assigned: the molecules written in bold belong directly to the shortened list, while the regular ones have been added to complete the pathway; the molecules written with the color that identifies the pathway (both bold and regular) have been validated by western blot analysis. **D)** Western blot validation of selected signaling molecules from the above mentioned signaling pathway during VSVG traffic pulse from the TGN to the PM. **E)** Western blot analysis of the dependence on the presence of the cargo in the TGN for the activation of the signaling during the temperature shift at 20°C in the presence or absence of Cycloheximide (CHX, 50ug/ml). The Cycloheximide was added at 37°C half an hour before the shift to 20°C and maintained until the end of the assay to completely empty the TGN and avoid the arrival of newly synthesized cargo.

It is plausible that the above signals might be generated by cargo traversing earlier stations of the secretory pathway (Cancino et al., 2014; Subramanian et al., 2019). To address this, we performed the protocol based on cargo accumulation specifically at the TGN by incubating cells at 20°C. We then validated the activation of key signaling proteins in the above cascades by immuno-blotting. PKD, ERK1/2 and eIF2α were robustly phosphorylated during cargo accumulation at the TGN (Figure 1D and 1E), while STAT was dephosphorylated (Figure 1D). As a further control, the same temperature shift in absence of cargo (cells treated with cycloheximide, Figure 1E) did not elicit any of the above-mentioned responses, indicating that the observed signaling response is exclusively dependent on the cargo accumulations and it is not a byproduct of the temperature changes or other manipulations.

We next asked if the activation of the above pathways might be responsible for cargo export from the TGN to the PM. We depleted or inhibited the relevant kinases/molecules and monitored the PM arrival of basoltareal cargo protein VSVG. Strikingly, we found that only the inhibition of PKD caused a block of VSVG exit from the TGN (data not shown).

This is in concordance with earlier studies that indicated the requirement of PKD activity for the formation of basolateral cargo carriers and their traffic to the PM (Yeaman et al., 2004). PKD is activated downstream of a signaling pathway that comprises Gβγ, PLCβ3, DAG, and PKC (Diaz Añel A.M. 2007; Lau et al. 2013; Diaz Añel and Malhotra, 2005). We confirmed that either the depletion of PLCβ3 or the inhibition of Gβγ significantly arrested cargo at the TGN (Supplementary Figure 1A and 1C).

These results in sum suggest that cargo accumulation at the TGN activates multiple signaling events. Specifically, the cargo dependent activation of PKD inturn regulates cargo export to the basolateral PM.

### Basolateral cargo overload in the TGN lumen triggers the activation and recruitment of PLCβ3 and PKD on the TGN membrane

The targeted phosphoproteomics analyses indicated an activation of PLC-PKD signaling as a consequence of cargo arrival at the TGN. We hence focused on examining the recruitment of these signaling components to TGN membranes in response to TGN cargo accumulation.

First, we determined which specific cargoes at the TGN elicited PKD activation. PKD is mainly cytosolic at steady state and is recruited to membranes by DAG via the kinase’s C1a domain (Maeda et al., 2001). Its activation can be detected by assessing its autophosphorylation at position ser916, a signal diagnostic of the kinase activity (Matthews et al., 1999). We co-transfected HeLa cells with GST-tagged PKD1 and with various synchronizable cargoes that traffic to either the basolateral PM (E-cadherin, TNFα and hGH) or the apical PM (GPI-GFP), and examined the effect of elevated TGN cargo loads on the recruitment and activation of PKD at the Golgi (Figure 2D and Supplementary Figure 2 A-B-C). The cargo proteins were first accumulated in the ER and then allowed to exit and accumulate in the TGN compartment at 20°C for 2 hours. Release of these cargoes from the TGN was then induced by shifting the cells to the permissive temperatures. Strikingly, the accumulation of only the basolateral cargoes in the TGN lumen resulted in the activation and recruitment of PKD to the TGN (Figure 2D). The apical cargo GPI-GFP had no effect (Supplementary Figure 2C).

**Figure 2:**
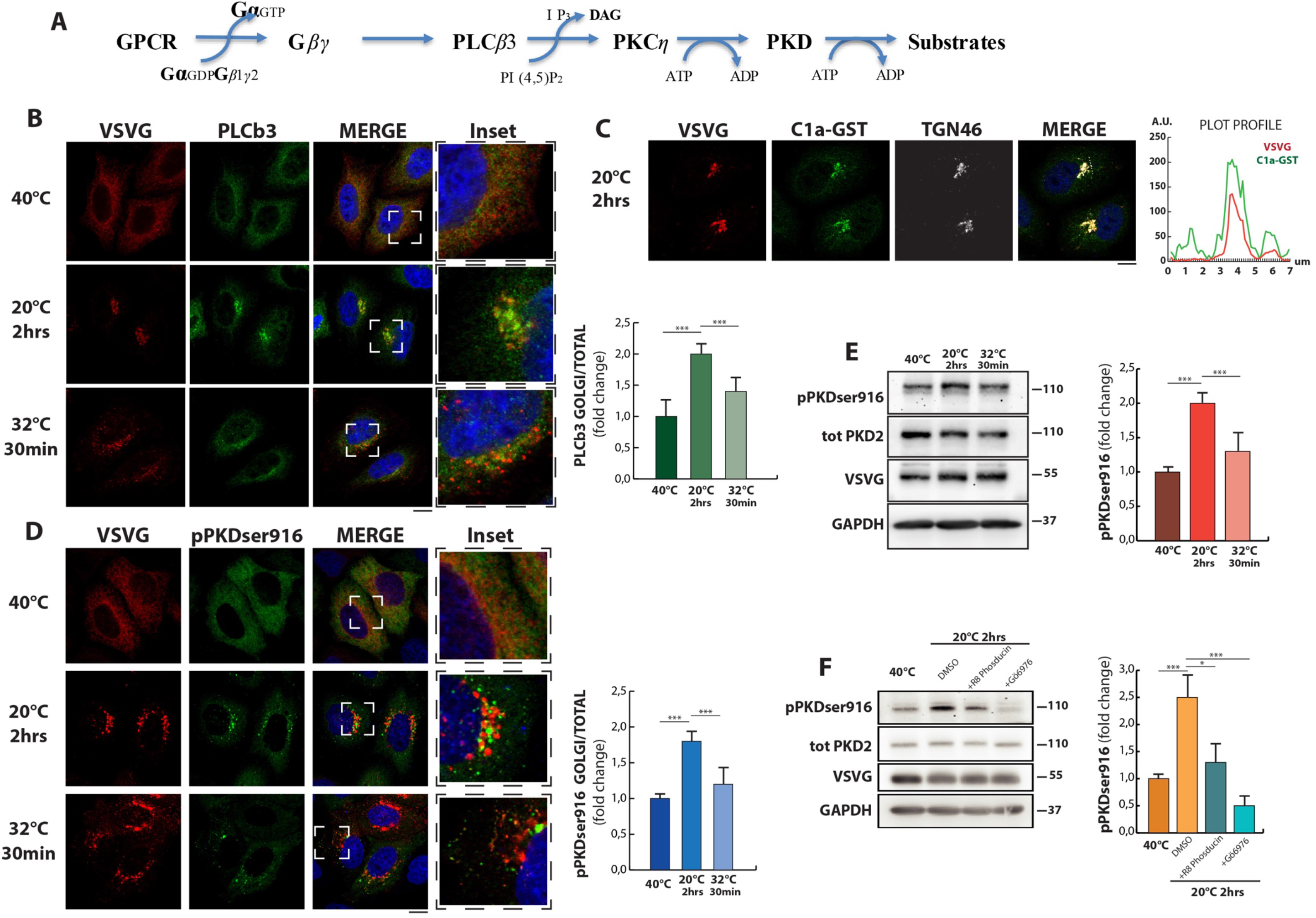
Basolateral cargo accumulation in the TGN induces a local recruitment and activation of the PLCb3-DAG-PKD signaling pathway. **A)** Conventional signaling cascade that leads to PKD activation on TGN membranes. **B)** IF analysis of the localization of the endogenous protein PLCb3 in HeLa cells during the VSVG traffic pulse at the indicated time points. The graph represents the PLCb3 levels in Golgi compared to the total. **C)** Localization of the DAG-reporter (C1a-PKD-GST) during VSVG accumulation in the TGN. The graph shows the co-localization profile between VSVG and the C1a-PKD-GST. **D)** IF analysis of the localization of the active form of the PKD protein by labeling the PKD1-GST overexpressed with the anti pPKDser916 antibody (auto-phosphorylation residue) during the traffic pulse of VSVG. The graph highlights the levels of pPKDser916 in the Golgi at the indicated time points. **E)** Western blot analysis of endogenous pPKDser916 levels during VSVG traffic pulse. **F)** Western blot analysis of endogenous pPKDser916 levels during VSVG traffic pulse at 40°C, and at the time point of the 20°C block in the presence of the beta-gamma subunit inhibitor (R8-phosducin aa215-233, 50uM, 1h, lane 3) or PKD (Gö6976 10uM, 1h, lane 4), or vehicle (DMSO, lane 2).

An essential step for PKD activation is the local DAG production that has a dual function of recruiting and activating PKD via PKC dependent phosphorylation (Maeda et al., 2001; Diaz Añel and Malhotra, 2005). A common approach to monitor DAG production in cells is to use the C1a-(PKD)-GST plasmid derived from the DAG binding domain of PKD, which is recruited to cell membranes enriched with DAG. To assay for DAG production on the TGN membranes, HeLa cells were transfected with the C1a-(PKD)-GST plasmid and a TGN cargo pulse was carried out. C1a-(PKD)-GST was recruited to the TGN membranes during TGN accumulation of VSVG at 20°C (Figure 2C).

The generation of DAG on TGN membranes during basolateral cargo secretion is possibly due to the action of the PLCβ3 (Diaz Añel, 2007; Sicart et al, 2015; Anitei et al,. 2017). PLCβ3 is a predominantly cytosolic protein that can be recruited to membranes by activation of Gβγ subunits (Fogg et al., 2001). PLCβ3 can also localize at the Golgi and is necessary for the TGN export of basolateral cargoes (Diaz Añel, 2007; Anitei et al,. 2017). We tested the effects of elevated TGN cargo levels on the distribution of PLCβ3. At steady state, PLCβ3 localized mostly in the cytosol and on the TGN (Figure 2A). During TGN accumulation of the cargo at 20°C, PLCβ3 markedly shifted from the cytosol to the Golgi. Then, as cargo left the TGN for the PM, upon release of the TGN 20°C temperature block, it redistributed back to the cytosol (Figure 2B). Since PLCβ3 is activated by the Gβγ subunit, we tested the effect of an inhibitory peptide of the Gβγ subunits (Blüml et al., 1997). Gβγ inhibition blocked both the cargo dependent PKD activation (Figure 2F) and the arrival of VSVG on the PM (Supplementary Figure 1C). These data overall indicate that the PLCβ3-PKD pathway activated by cargo at the TGN, in accordance with the activation of a pathway previously reported to be necessary for the TGN export of basolateral cargo (Malhotra and Campelo, 2011).

Altogether, these data suggest the presence of an autoregulatory mechanism that senses the amount of cargo in the TGN lumen and triggers a Gβγ-PLCβ3-PKD pathway that activates cargo export out of the TGN. We named this autoregulatory system as ARTG (**<u>a</u>**uto**<u>r</u>**egolatory system for **<u>TG</u>**N export).

### Basolateral cargo export from the TGN requires the orphan receptor GPRC5A

The core of an autoregulatory circuit is the presence of a sensor that detects the critical variable to be controlled, which in this case is the load of basolateral cargo at the TGN. For the ARTG system, we predicted that a sensor must be situated on the Golgi membrane. To identify the putative sensor of TGN cargo load, we resorted to an unbiased ‘interactomics’ approach based on the assumption that the sensor might be directly or indirectly associated with the accumulating cargo. We immunoprecipitated VSVG at three distinct time points during a cargo transport pulse: i) at 40°C when VSVG is in the ER; ii) at 20°C when VSVG accumulates at the TGN; and iii) 30 minutes after release from the TGN at 32°C when VSVG actively traffics in carriers destined toward the basolateral PM. Under each of these conditions, the cargo interacting partners were detected by iterative rounds of mass spectrometry (Figure 3A). Strict filtering based on a ranked score (see Figure 3B) resulted in a list of approximately 100 interacting proteins that were subsequently analyzed. We focused on cargo interacting partners that were present only during TGN accumulation and after cargo release from the TGN, and excluded those proteins that interacted with cargo arrested in the ER (Figure 3B). Remarkably, our proteomics screen highlighted an interaction between cargo and a member of the class C of GPCRs, named GPRC5A (Cheng and Lotan, 1998; Ye et al., 2009). Co-immuno precipitation and immuno-blotting confirmed that GPRC5A is a *bona fide* interactor of basolteral cargoes VSVG and hGH at the TGN (Figure 3C and 3D), suggesting that this orphan receptor might initiate the signaling events that activate the Gβγ-PKD pathway.

**Figure 3:**
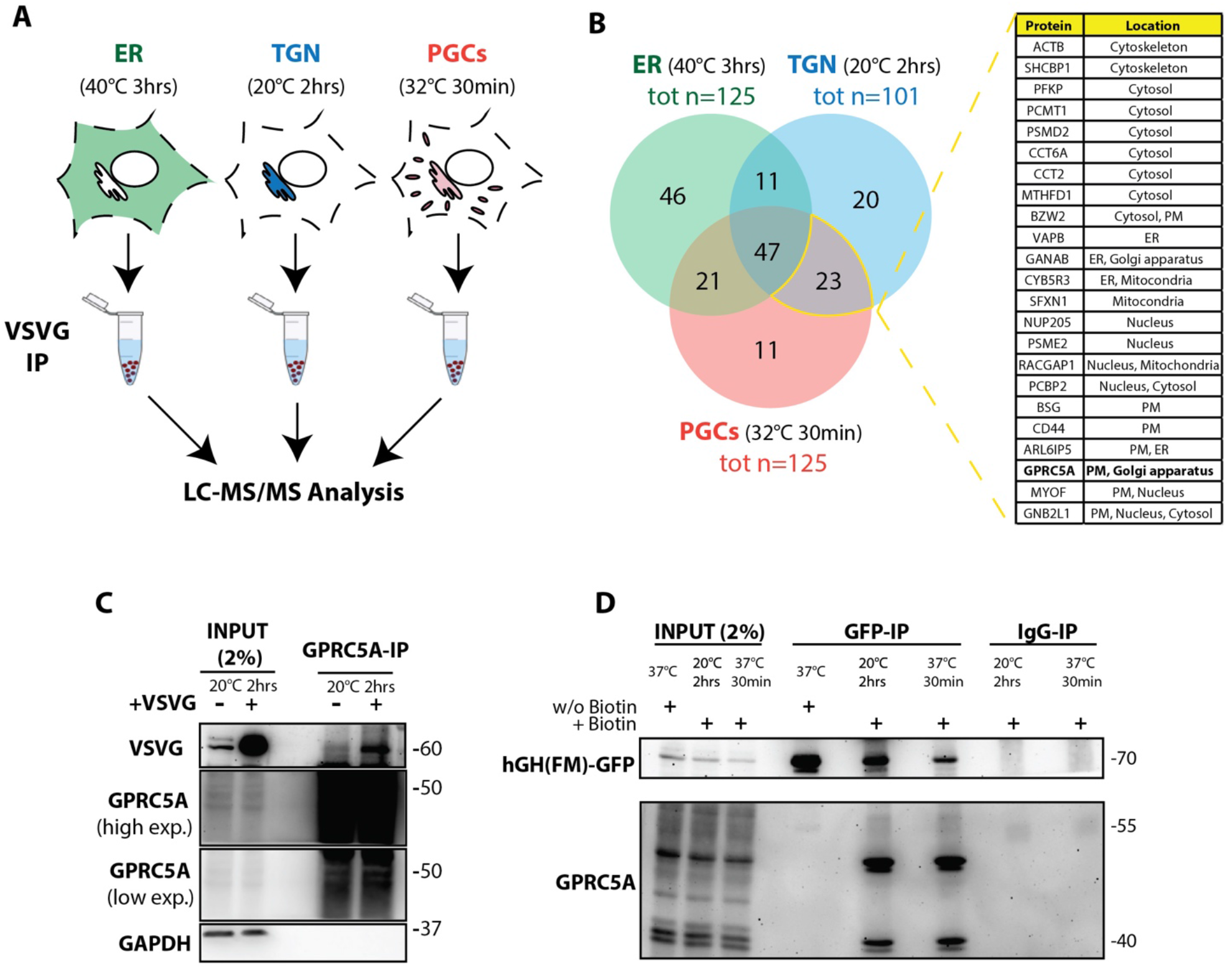
Proteomic analysis of VSVG interactome in the different stations of the secretory pathway reveals GPRC5A as novel cargo interactor in the TGN compartment. **A)** Experimental design of the VSVG binding proteins analysis across the secretory pathway. **B)** Venn diagram of VSVG putative interactors in the different samples. We applied some filters to exclude un-specifically bound proteins using protein score between 500 and 120 (assigned by the mass spec facility as the sum of the single peptide scores) as arbitrary cut-off. From this step of analysis, we excluded the proteins in common to all the samples, and then we focused on the proteins found to co-immunoprecipitate with VSVG both in the TGN and in the PGCs (samples 20°C and 30 minutes at 32°C). Those proteins are listed in the table along with their subcellular localization. **C)** Westen blot analysis confirms that GPRC5A co-immunoprecipitates with VSVG during the 20C block. **D)** Co-IP analysis of the interaction between the soluble basolateral cargo hGH and GPRC5A during hGH traffic pulse.

GPRC5A localizes to the PM, Golgi and post Golgi vesicles (Zhou and Rigoutsos, 2014). We confirmed that the endogenous protein localizes to the Golgi and perinuclear vesicles in multiple cell types (Supplementary Figure 4A). We then investigated the potential role of this GPCR in the TGN export of cargoes. GPRC5A depletion resulted in a significant block of basolateral cargoes VSVG, E-cadherin, TNFα, and hGH exiting the TGN (Figure 4A, Supplementary Figure 4C-E). An siRNA resistant tagged-mutant GPRC5A when expressed rescued cargo transport to the PM (Figure 4D). Importantly, GPRC5A depletion did not affect the export of apical cargoes GPI-GFP and GP135-GFP and lysosomal cargoes LAMP1, Cathepsin D and heparanase (Supplementary Figure 4B-D-F and data not shown).

**Figure 4:**
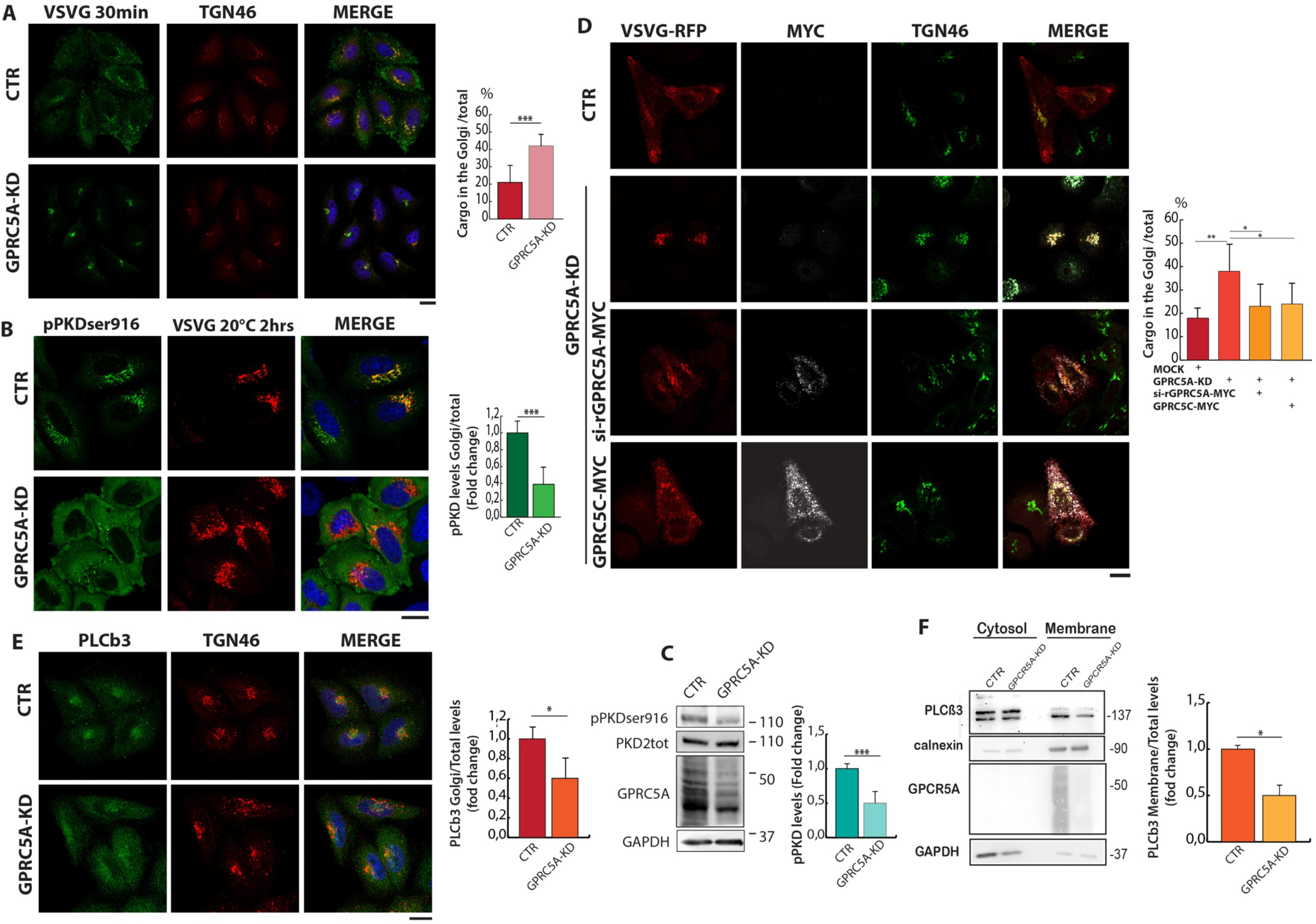
GPRC5A regulates through the pathway that leads to the activation and recruitment of the protein Kinase D. **A)** IF analysis of VSVG protein transport at 30min time at 32°C (after block at 20°C for 2 hours) in HeLa cells. The quantification in the graph shows the accumulation of VSVG in the TGN in the GPRC5A-KD cells compared to the control cells. **B)** IF analysis of the localization of the active form of the protein kinase D (overexpressed PKD1-GST and labeled with the anti pPKDser916 antibody) in control cells and GPRC5A-KD during the traffic pulse with VSVG in the block time point at 20°C. The quantification shows that in the GPRC5A-KD cells the active form of PKD is not recruited to the TGN with respect to the control cells. **C)** Rescue experiment of the transport of VSVG in GPRC5A-KD cells by expression of the GPRC5A siRNA resistant construct (si-rGPRC5A-Myc) or of the GPRC5(B-C-D)-Myc constructs. The IF analysis of VSVG transport shows that both si-rGPRC5A-Myc and GPRC5C-Myc constructs are able to recover VSVG traffic defects induced by the depletion of GPRC5A, while the overexpression of GPRC5B or GPRC5D does not recover (data not shown), indicating a potential partial redundancy of function of the proteins GPRC5A and GPRC5C. **D)** Western blot analysis of pPKDser916 levels during VSVG traffic pulse in control and GPRC5A-KD cells. **E)** IF analysis of endogenous PLCb3 localization in control and GPRC5A-KD HeLa cells. Quantification in the graph highlights a reduced Golgi localization of PLCb3 in GPRC5A-KD cells compared to control cells (scale bars 10nm). **F)** Western blot analysis of cytosolic and membrane fractions of control and GPRC5A-KD steady state HeLa cells. The quantification in the graph shows the ratio of the membrane fraction of PLCb3 to the total using Calnexin (membrane fraction) and GAPDH (cytosolic fraction) as normalizers.

These results strongly indicate a specific role for GPRC5A in the transport of basolateral cargoes, and hence the regulation of sorting stringency at the TGN.

### The ARTG-PKD pathway is regulated by GPRC5A

Our results so far suggest that GPRC5A senses basolateral cargo arrival at the TGN and subsequently activates downstream heterotrimeric G proteins, thus initiating the signaling cascade that culminates with PKD activation. To test this model, we depleted GPRC5A and assayed for PKD activation. Indeed, we observed a decrease in the activated form of the PKD (Ser916) by immuno-blotting (Figure 4C). Moreover, GPRC5A depleted cells exhibited a poor recruitment of PKDser916 and PLCβ3 to the TGN and a predominant cytosolic localization, even under conditions of substantial TGN cargo accumulation (Figure 4B and 4E). At steady state (i.e.) in the absence of cargo overloads, PLCβ3 was poorly recruited to the membrane fraction in cells depleted of GPRC5A (Figure 4F).

As stated previously, both PLCβ3 and PKD activation requires Gβγ subunits (Diaz-Añel and Malhotra, 2005). Classically, G-protein coupled receptors (GPCR) remain in complex with heterotrimeric Gα and Gβγ subunits in an inactive state (Hilger et al., 2018). GPCR activation results in a GDP to GTP exchange on the Gα subunit and a subsequent release of active Gβγ subunits to initiate subsequent downstream signaling events (Hilger et al., 2018). To identify the Gα protein(s) activated by the GPRC5A during TGN cargo accumulation, we performed a second unbiased ‘interactome’ screen by pulling down GPRC5A and analyzing the interacting partner proteins by mass spectrometry (Figure 5A). We identified the heterotrimeric G-proteins Gαi3, Gβ2 and Gγ12 in complex with GPRC5A and determined Gαi3 as a *bonafide* interactor of GPRC5A by immunoblotting (Figure 5B).

**Figure 5:**
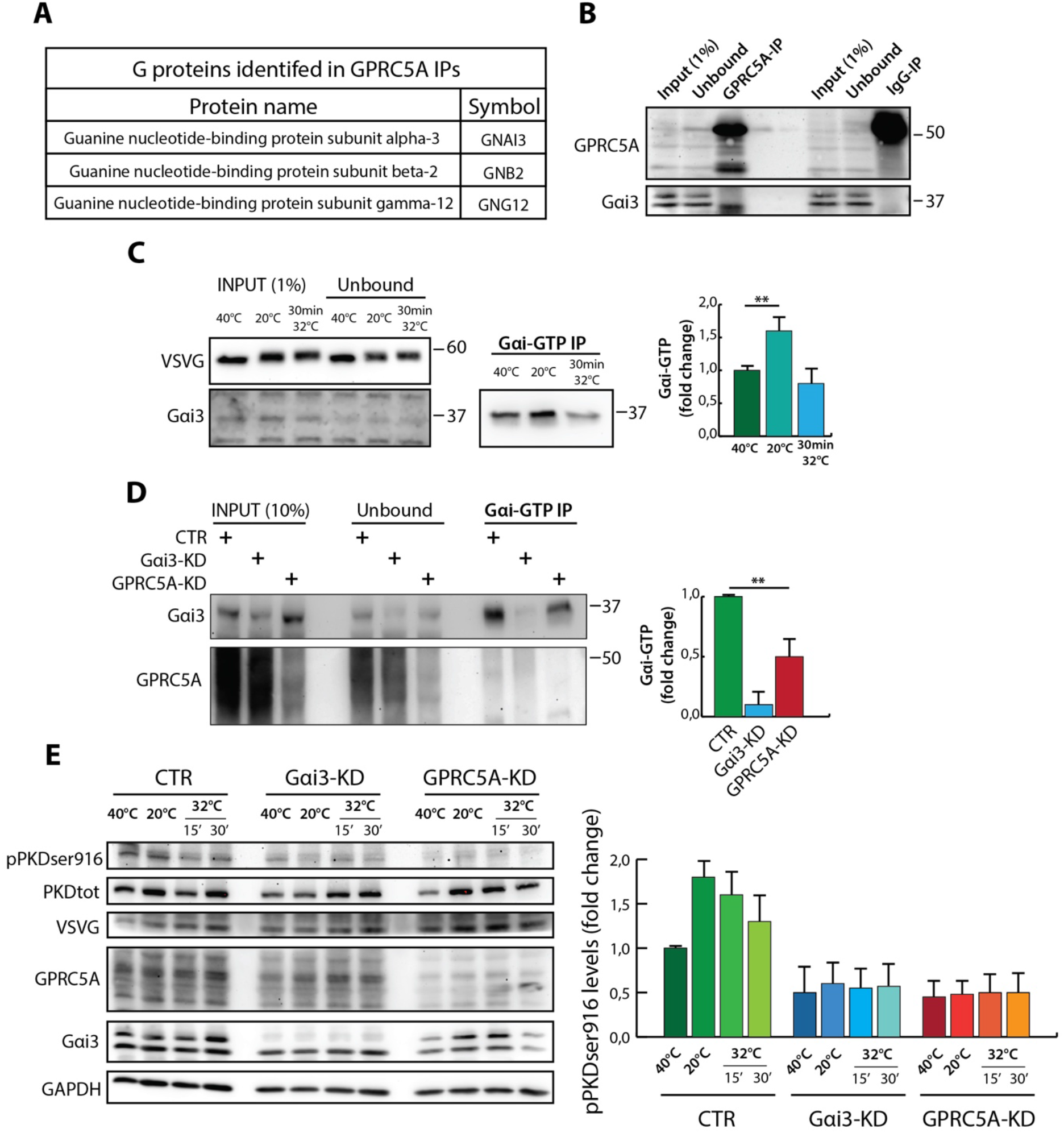
G alpha i-3 is coupled to GPRC5A and it is activated during cargo accumulation in the TGN. **A)** List of G proteins identified in GPRC5A IPs by LC-MS/MS analysis. **B)** Western blot (WB) validation of Gαi-3 co-immunoprecipitation with GPRC5A in steady state HeLa cells. **C)** WB analysis of Gαi-3 active form (Gαi-GTP) immunoprecitiation during VSVG traffic pulse. The quantification highlights the increased amount of active Gαi-3 during VSVG accumulation in the TGN. **D)** WB analysis Gαi-GTP inmmunoprecipitated levels in control, GPRC5A-KD and Gαi-3-KD steady state HeLa cells. The quantification shows the decreased levels of active Gαi-3 in absence of GPRC5A, Gαi-3-KD cells are used as negative control of the assay. **E)** WB analysis of PKDser916 levels in control, GPRC5A-KD and Gαi-3-KD during VSVG traffic pulse in HeLa cells. The graph clearly shows that in either GPRC5A-KD or Gαi-3-KD condition there is no activation of PKD in response to VSVG accumulation in the TGN compartment.

We then investigated if Gαi3 is activated during cargo accumulation at the TGN. We measured the amount of activated Gαi3 (Gαi3-GTP) by taking advantage of a characterized antibody that recognizes and immunoprecipitates the GTP-bound (see Methods for details). Immunoblotting analyses of immunoprecipitated Gαi3 showed a significant increase in the active GTP bound form of Gαi3 only when cargo accumulated at the TGN (Figure 5C). Depleting GPRC5A in steady state HeLa cells resulted in a decrease of active Gαi3 compared to controls (Figure 5D). Significantly, GPRC5A and active Gαi3 were important for cargo induced PKD activation at the TGN as their absence resulted in a complete loss of PKD phosphorylation (Figure 5E), suggesting that the two proteins act in the same signaling pathway.

Finally, we also tested if GPRC5A signaling regulated the intensity the eIF2α phosphorylation. Strkingkly, eIF2α phosphorylation was markedly increased in GPRC5A-KD cells both at steady state and during cargo overload at the TGN (data not shown), presumably as a consequence of aberrant TGN cargo accumulation. The signaling events that result in this marked phosphorylation remain to be understood.

Altogether these results suggest that the ARTG signaling pathway on TGN membranes is initiated by basolateral cargo interactions with GPRC5A. The cargo-GPRC5A complex activates Gαi3 and the release of Gβγ subunits, that in turn activate the PLCβ3-nPKC-PKD pathway. PKD activation induces the export of basolateral cargoes from the TGN. Notably, ARTG does not control the export of apical or lysosomal cargoes from the TGN, highlighting the possibility that this signaling circuit maintains optimal sorting and polarity in cells.

### Mechanism of ARTG activation by basolateral cargo through GPRC5A

Our data suggest that basolateral cargoes in the TGN lumen interact with GPRC5A, or with a protein complex containing GPRC5A. The analysis of this interaction in the lumen of the TGN is complicated by an abundance of factors (pH, ionic concentration, other types of proteins and complex carbohydrate and lipid composition) (Paroutis et al., 2004; Boutté Y., 2018; Kellokumpu, 2019).

To circumvent these issues, we sought to study the interaction between basolateral cargoes and GPRC5A under simplified conditions by expressing the receptor on the cell surface. This is possible since overexpression of GPRC5A results in extensive PM localization, in addition to its Golgi and PGC localization (Figure 6F). Similar results are also achieved by treating cells with retinoic acid (RA) (Figure 6B), which is known to increase the expression of the receptor (Cheng and Lotan, 1998; Ye et al, 2009). Under these conditions, which results in a homogenous localization of the receptor at the PM, cells were first washed with buffer and then incubated with soluble basolateral cargoes, such as TNFα and albumin, or the apical cargo, lysozyme. Soluble cargo proteins were added in serum-free media for 15 minutes to the cells pre-incubated at 20°C for 3 hours to prevent further transport of GPRC5A to or from the plasma membrane. The soluble basolateral cargoes (albumin or TNFα) induced a marked activation of PKD (Figure 6C and 6D) and recruitment of the activated kinase to the PM (Figure 6F), while the soluble apical cargo lysozyme had no effect (Figure 6A). This differs from the effects of cargo on GPRC5A, which results in PKD recruitment at the TGN. In control cells that do not express GPRC5A at the PM, the addition of albumin or TNFα to the cell medium had no effect on PKD. A schematic model of the results from these experiments is described in Figure 6H. The dose-response curves for these cargoes identify an EC50 of 0.28 mM for Albumin and 1.3 nM for TNFα (Figure 6G). These values are compatible with the high secreted levels of albumin and the low levels for TNFα considering that the affinity of these proteins for their own receptors is in the order of 100pM for TNFαR (David J. MacEwan, 2002) and in the order of 6-600 nM for different types Albumin receptors (Bern et al., 2015).

**Figure 6:**
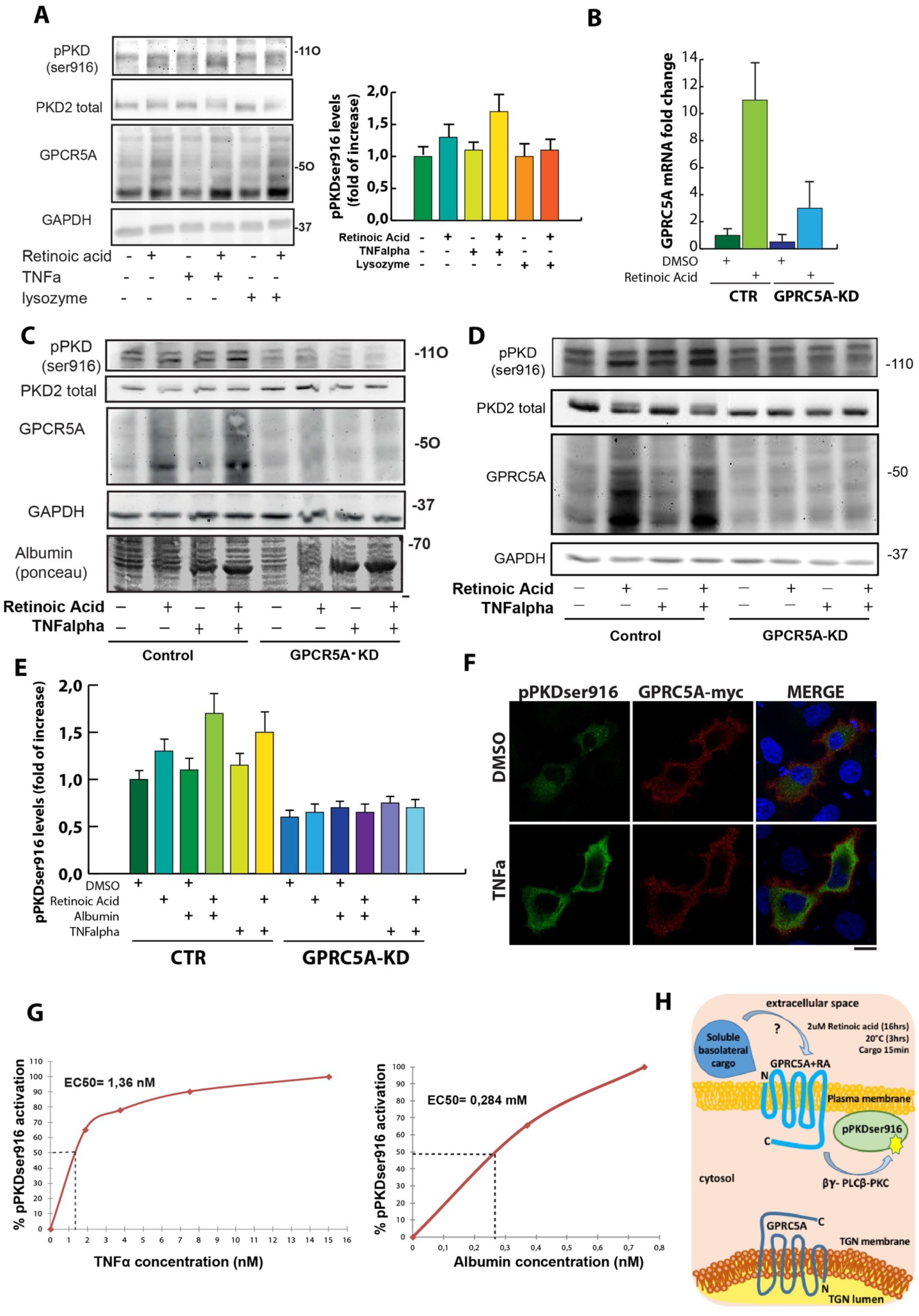
GPRC5A on plasma membrane can activate PKD in response to the addition of soluble cargo in the extracellular medium. **A)** RT-PCR analysis of GPCR5A transcript levels upon DMSO or retinoic acid (RA) treatment (2uM, 16hrs in serum-free media) in control and GPRC5A-KD HeLa cells. **B)** Western blot analysis of endogenous pPKDser916 levels in cells treated with RA (or DMSO) and two different soluble cargoes, TNFα (15nM, basolaterally secreted) and Lysozyme (15mM, apically secreted), or vehicle. After RA (or DMSO) treatment all conditions have been incubated at 20°C for 3hrs in serum-free media, then soluble cargoes have been added for 15 minutes. **C-D)** Western blot analysis of endogenous pPKDser916 levels in control (lanes 1 to 4) and GPRC5A-KD (lanes 5 to 8) cells treated with RA (2uM, 16hrs in serum-free media, lanes 2,4,6 and 8) and albumin (750uM, 15min, panel Clanes 3,4,7 and 8) or recombinant TNFα (15nM, 15min, panel D lanes 3,4,7 and 8). All conditions have been performed as above mentioned for panel B. **E)** Quantification of pPKDser916 levels from C and D experiments. **F)** IF analysis of pPKDser916 (overexpressed PKD1-GST) localization in cells overexpressing GPRC5A treated with recombinant TNFα or vehicle (scale bar 10nm). **G)** Dose-response curves for pPKDser916 activation with different concentration of TNFα and albumin. **H)** Graphical summary and model of the experiments described in this figure.

Notably, the addition of cargo to the above-described system at 37°C did not activate PKD. From a physiological point of view, the absence of receptor activation at the temperature of 37°C could mean that under basal conditions this signaling machinery cannot be erroneously activated by the soluble basolateral cargoes that are normally present in the extracellular space.

### Receptors of the GPRC5 family have a partially redundant function in different cell lines

We expect the ARTG to be a basally active in all cells, even though the specific members of the GPRC5 family might differ in expression levels. GPRC5 family members are differentially expressed among healthy human tissues (Zhou and Rigoutsos, 2014). We asked if any of the other GPRC5 paralogues replace GPRC5A function, thus implying redundant roles in TGN export. The GPRC5 family members in mammals are GPRC5A, GPRC5B, GPRC5C, and GPRC5D; with all members being orphan receptors (Bjarnadottir et al, 2005). GPRC5A is expressed at high levels in the lung (Tao et al., 2004; Xu et al., 2005), whereas the expression pattern of GPRC5B is mostly restricted to neurons with limited expression in other tissues (Brauner-Osborne et al., 2000; Robbins et al., 2000; Robbins et al., 2002; Kim et al., 2018). The expression of GPRC5C is more widespread than GPRC5A/B and is prevalent in the kidney and liver (Robbins et al., 2000). GPRC5D is associated with hard-keratinized structures, like cortical cells of the hair shaft (Inoue et al., 2004). We ectopically expressed myc tagged versions of the isoforms in GPRC5A depleted HeLa cells and assayed for cargo transport to the PM. As shown in Figure 4D, only GPRC5C was able to rescue VSVG trafficking out of the TGN, while the other two GPCRs had no effect (data not shown). GPRC5A and GPRC5C share around 30% amino acid identity, but localize to similar compartments in HeLa cells (Supplementary Figure 4B), supporting the hypothesis that GPRC5C could have redundant functions to GPRC5A in other cell types, such as hepatocytes, where it is majorly expressed.

### Role of the ARTG pathway in polarized secretion and cell polarity

We finally explored the functions of the ARTG in regulating cell polarity with focused experiments in polarized HepG2 hepatocytes. This cell line has been extensively used to study membrane polarity (Anne Müsch, 2014) and expresses high levels of GPRC5C. In addition, hepatocytes are professional secretory cells presumably high burdens of cargo proteins (such as albumin) traversing the secretory pathway. Normal polarized HepG2 show different cell membrane domains where the asymmetry is achieved through the establishment of tight junctions that mark the borders between apical and basolateral membranes such that apical domains of adjacent hepatocytes form the bile canaliculi whose function is to excrete bile salts from the cell (Gissen and Arias, 2015). Bile canaliculi formation is an important feature of polarized hepatocytes and can be observed as channels that are clearly marked by a dense bundle of actin microfilaments and a ring-like structure of the tight junction zonula occludens 1 (ZO-1), as shown in Figure 7D.

**Figure 7:**
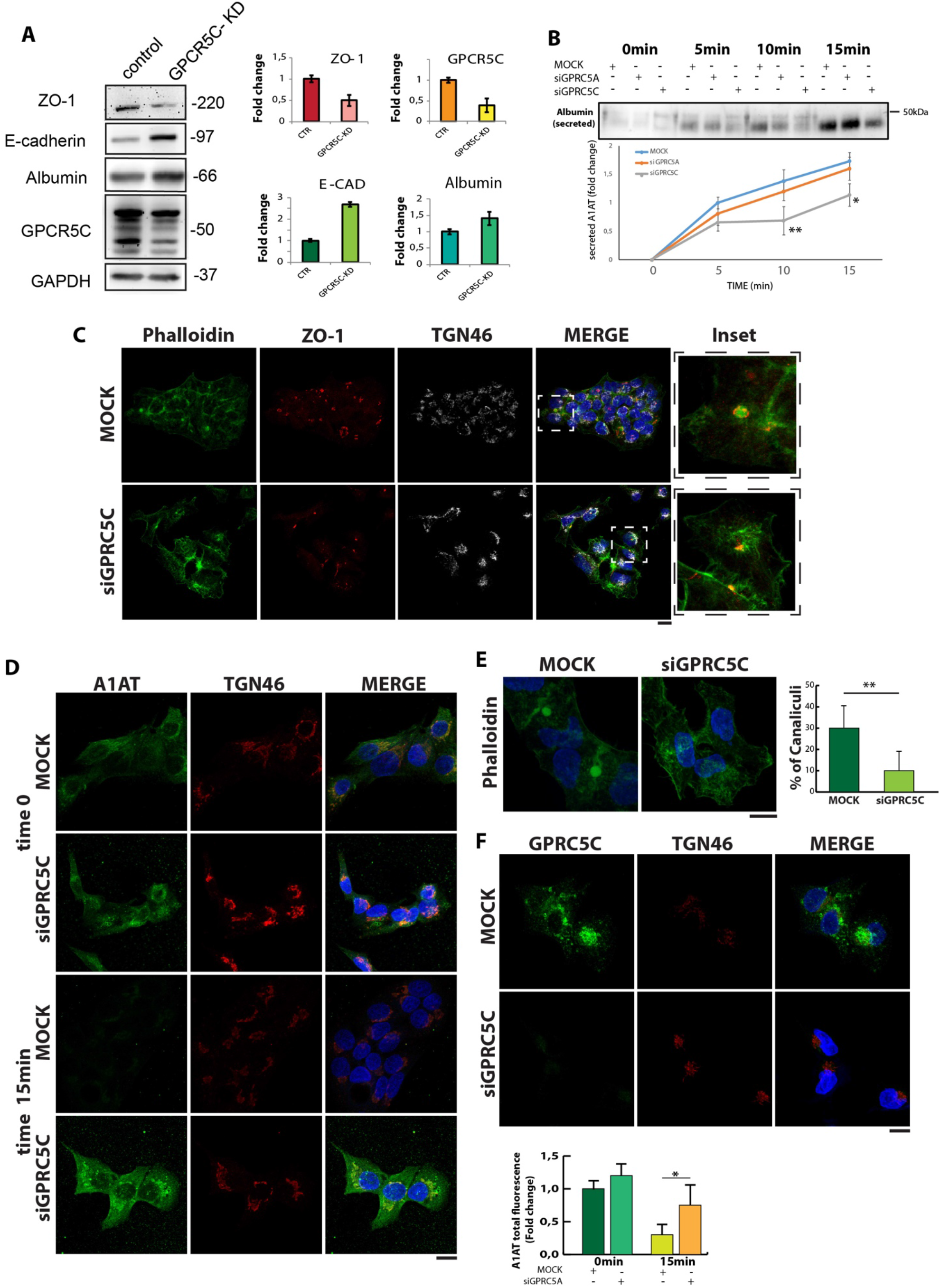
GPRC5C regulates polarization and secretion of basolateral soluble proteins in HepG2 cells. **A)** IF analysis of the localization of the endogenous alpha1 anti-trypsin protein (A1AT) in control and GPRC5C-KD HepG2 cells at 20°C for 2hrs (time 0) and 15 minutes after the shift of temperature to 37°C. **B)** Western blot analysis of A1AT secreted in the medium of HepG2 control and GPRC5A-KD cells at different time points at 37°C after blocking at 20°C for 2 hours (time 0). **C)** IF analysis of Phalloidin-488 staining to visualize the bile-canaliculi, visualized as a circular and dense structure of variable diameter, in steady state control and GPRC5A-KD HepG2cells. The graph represents the percentage (%) of canaliculi in both conditions. **D)** IF analysis of the localization of the junction protein Zonula occludens 1 (ZO-1), in control and GPRC5C-KD cells. Co-staining with Phalloidin shows that in control cells the ZO-1 protein forms a ring around the actin that forms the canaliculus, while in GPRC5C-KD this does not happen and ZO-1 remains localized inside the cell or to the cell periphery (scale bars 10nm). **E)** Western blot analysis of different polarity and cell junction markers in control and GPRC5C-KD HepG2 cells, with their relative quantification. **F)** IF analysis of GPRC5C in control and GPRC5C-KD HepG2 cells.

**Figure 8:**
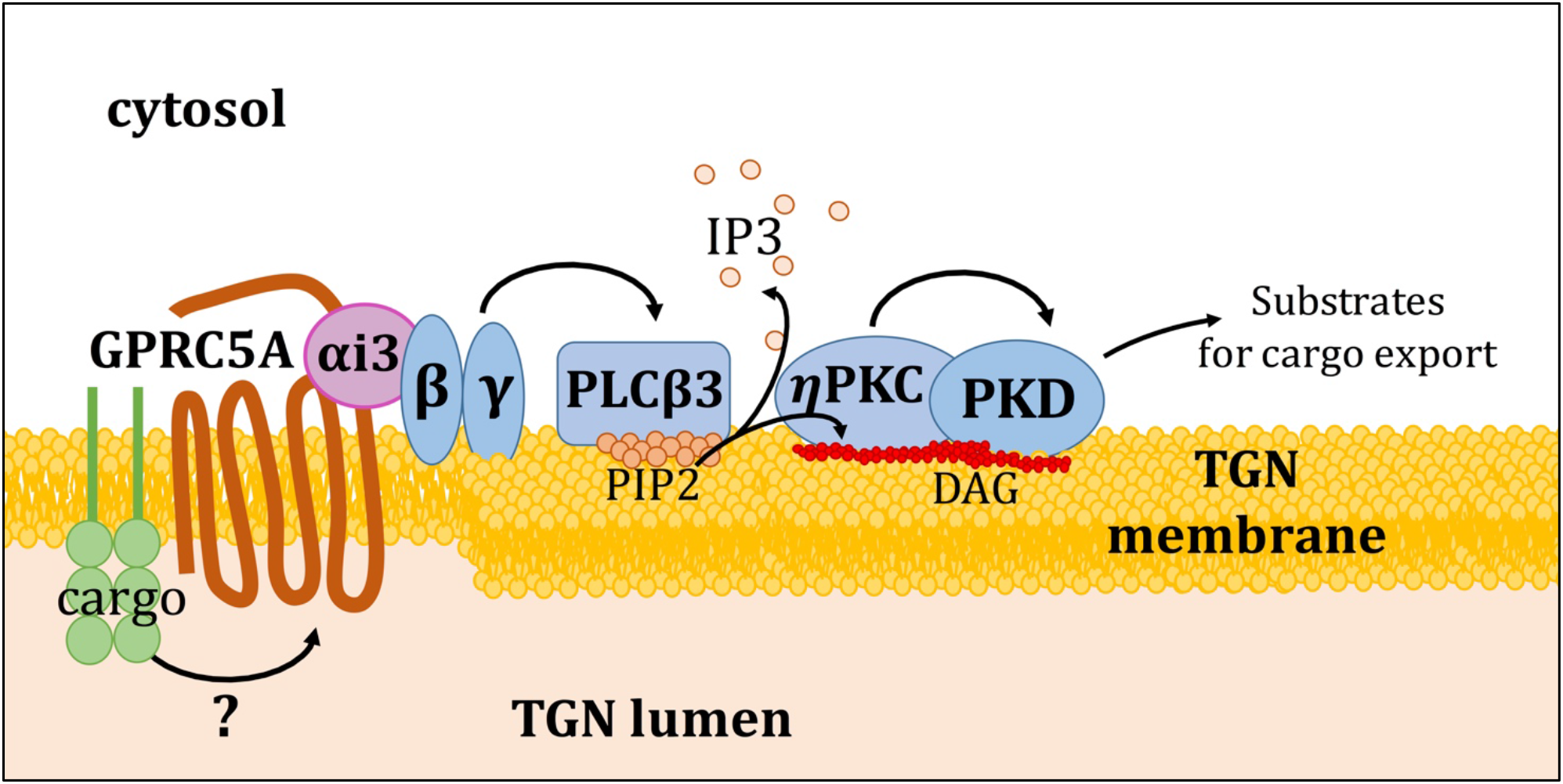
Cargo dependent ARTG signaling activation model. Basolateral cargoes arrival in the TGN, from which they must be subsequently sorted, activate the orphan receptor GPRC5A. The mechanism that directly associates the cargo to the receptor is still being analyzed, however, the associated Gβγ subunits start a signaling cascade that flows onto the activation and recruitment of PKD. Previous literature had, in fact, identified the Gβγ subunits, PLCβ3 and ηPKC as molecules upstream of the activation of PKD on the TGN (Jamora et al., 1999; Baron and Malhotra, 2002; Diaz Añel and Malhotra, 2005; Diaz Añel, 2007). The final effect of this signaling is the phosphorylation by PKD of substrater required for the export of basolateral cargoes. We called this circuit ARTG, for **A**uto**r**egulation of **TG**N export.

We examined the effects of GPRC5C depletion on the trafficking of endogenous basolateral proteins, such as ZO-1, E-cadherin, and albumin in HepG2 cells (Figure 7A). Both immunoblotting (Figure 7A) and microscopy (Figure 7C) analyses revealed gross alterations in the polarity marker ZO-1. Moreover, GPRC5C-KD cells did not form clusters with an apparent loss of a polarized PM domain architecture. Interestingly, the levels of E-cadherin were increased, a phenotype linked with a disrupted hepatic cell polarity (Konopka et al., 2007).

We next investigated the effects of GPRC5C depletion on bile canaliculi formation. Usually, around 30% to 40% of HepG2 cells (cultured in a monolayer for at least 72hrs) polarize with bile canaliculi-like structures (Van IJzendoorn et al., 2004). GPRC5C depletion resulted in a dramatic reduction in the percentage of cells with bile canaliculi (~7%; Figure 7E). Moreover, intracellular albumin (Figure 7A) and alpha 1 anti-trypsin (Figure 7D) levels were elevated in GPRC5C depleted cells, suggesting defects in the normal secretion of these proteins. Indeed, the secretion of albumin into the extracellular medium was markedly delayed in cells depleted of GPRC5C (Figure 7B).

These results indicate a role for ARTG in the sorting and secretion of endogenous soluble basolateral cargoes and in the optimal polarization of the HepG2 cells.

## Discussion

In this study, we report that the TGN is equipped with an autoregulatory molecular device, the ARTG complex, that is activated by the binding of basolateral cargo to an orphan receptor, GPRC5A (also known as RAI3) in the TGN and then activates the TGN basolateral export machinery. ARTG can thus be modeled as a control device operating to keep the basolateral cargo levels at low levels to avoid the potentially harmful effects of cargo overloads.

GPRC5A acts as a cargo sensor and signal transducer and, upon binding (directly or indirectly) with basolateral cargo proteins, induces the recruitment and activation of a set of signaling components that include the G protein Gαi3 and Gβγ subunits, PLCβ3, the production of DAG, and the phosphorylation of PKD. The activation of PKD results in the formation of a post-Golgi carrier targeted to the basolateral membrane. Several components of this pathway, from the Gβγ subunits to PKD had been identified previously. Their activation was proposed to be initiated at the PM by the stimulation of a GPCR-G proteins complex and the consequent release and translocation to the TGN of the Gβγ subunits (Diaz Añel, 2007). A different but analogous mechanism of activation of PKD at the TGN by PM receptors has been proposed by other authors (Eisler et al., 2018). Also Gαi3 has been previously reported to localize in the Golgi in different cell types (Stow et al., 1991a; Wilson et al., 1994) and to play role in intra-Golgi transport of heparan sulfate proteoglycan (Stow et al., 1991b). Moreover, recently, it has has been shown to be activated by a non-GPCR dependent mechanism at the Golgi (Lo et al., 2015). Gαi3 is also activated on other subcellular membranes, like LC3-positive autophagosomes (Garcia-Marcos et al., 2011), melanosomes (Young et al., 2011) and at the plasma membrane (Thompson et al., 2007; Kuwano et al., 2016). Thus, the same signaling molecules can operate in different ways in different regulatory networks for different purposes. The current study shows that the PKD pathway can be activated by cargo binding to GPRC5A-Gαi-3 at the TGN, and that it is a component of a regulatory device that supports optimal TGN functionality as well as cell polarity.

A mechanistic question raised by these findings concerns the recognition by GPRC5A of a multitude of basolateral cargoes, whose structural characteristics vary considerably. Sequence homologies and/or structural motifs common to all basolateral cargoes that would bind GPRC5A have not so far been identified by us or others. We propose a model in which different classes of basolateral cargoes activate the the GPRC5A receptor through one or more adaptors endowed with multiple binding sites, that form a complex with the GPRC5a receptor. This mechanism would be analogous to that proposed for the export of the large number of different protein species from the ER through multiple recognition domains in different Sec24 adaptors (Miller et al., 2003; Wendeler et al., 2007; Zanetti et al., 2012).

Related to the abve is the problem of the precise role of the GPRC5A complex in the sorting mechanism at TGN. The activation of GPRC5A by basolateral cargo proteins is essential for the selective transport of these proteins to the basolateral domain of the PM. One may however ask if the function of the GPRC5A complex is only due to the activation of the formation of basolateral carriers in which cargo proteins are then recruited by traditional sorting mechanisms, for instance by means of the clathrin adaptor (AP) complexes (Rodriguez-Boulan et al., 2005; Rodriguez-Boulan and Musch, 2005), or if the GPRC5A complex, which recognizes a lumenal region of the cargo to be activated, may also function as an adaptor for the recruitment of cargo in the basolateral carrier. The fact that only a fraction of the basolateral cargoes exhibit motifs that can be linked to known sorting mechanism (Di Martino et al., 2019) makes a direct role of the GPRC5A complex an attractive hypothesis, and in general the role of this complex in TGN sorting a fascinating territory to explore in this area.

Not all human tissues express GPRC5A. There are four mammalian genes belonging to the GPRC5 family, each with a specific expression profile across different human tissues. The other genes GPRC5 genes are GPRC5B, GPRC5C, and GPRC5D. At least some of them are involved in regulation of of basolateral cargo export in different cell types. GPRC5C manages to compensate for the lack of GPRC5A in restoring transport of basolateral cargo molecules (Figure 4D). Furthermore, GPRC5C, which is expressed at high levels in various tissues including the liver, regulates the secretion in HepG2 cells of endogenous basolateral cargo molecules, such as albumin, a1AT, and E-cadherin, and the consequent formation of cell junctions that determine the correct polarization of this particular cell type (Figure 7).

Many observations have highlighted a strong connection between alterations in GPRC5A and several human cancers (Jiang et al., 2018), and in particular with the presence of more aggressive and metastatic phenotypes through the induction of the EMT (Liu et al., 2016; Sawada et al., 2020; Liu et al., 2017).

EMT takes place physiologically during embryonic development, tissue regeneration, organ fibrosis, and wound healing. Through this process, epithelial cells can switch to a mesenchymal phenotype. This phenomenon is also involved in tumor progression with metastatic expansion and in the generation of cancer cells with stem cell properties, which play an important role in resistance to cancer treatment (Nieto et al. 2016, Lambert et al. 2017). E-cadherin is one of the main regulators of the EMT process. It is thought to act by maintaining cell-cell adhesion, with a global anti-proliferative, anti-metastatic, and anti-invasion effect. Epithelial polarity loss is a component of the EMT process, and plays a crucial role in the correct localization of key basolateral molecules, such as E-cadherin. GPRC5A depletion in mouse epithelial tracheal cells has been shown to cause a strong EMT induction following exposure to silica (Wang et al., 2015), suggesting a role for GPRC5A in maintaining epithelial identity especially in the context of lung tumors in which GPRC5A has been shown to act as a tumor suppressor (Tao et al., 2007). The current study indicates that the role of GPRC5A in cancer could be explained through the promotion and regulation of the mechanisms that support the activation of the machinery necessary for the export of basolateral molecules in particular by acting on PKD.

In MDCK cells, a polarized cell system (Yeaman et al. 2004), it has been found that both PKD1 and PKD2 play a critical role in trafficking of endogenous proteins to the basolateral membrane, such as E-cadherin and β1-integrin. By regulating the production of TGN carriers destined for the basolateral membrane, PKD1 and PKD2 can play an important role in the generation of epithelial polarity. In addition, PKD regulation has also been linked to cancer invasiveness through EMT. Evidence of reduced E-cadherin and PKD1 in advanced prostate, breast, and stomach cancers further supports this type of role for both proteins (Durand et al., 2016; Roy et al, 2017). The relationship between the two proteins is highlighted by several observations: PKD1 is essential to maintain the transcription of the E-cadherin gene and repress the expression of mesenchymal proteins through the phosphorylation of the transcription factor SNAIL (Du et al., 2010); activated PKD1 phosphorylates the cytoplasmic tail of E-cadherin, thus stabilizing its association with β-catenin and the actin cytoskeleton (Jaggi et al., 2005), which strengthens adherent junctions with consequent inhibition of cell motility. Therefore, the downregulation of PKD1 could facilitate the EMT phenomenon by compromising both E-cad function and expression and promoting the expression of genes involved in the mesenchymal phenotype.

The components of the ARTG machinery are also involved in genetic diseases and pathological conditions that can be related to defects in the correct protein secretion or cell polarization. Gαi-3 is associated with a rare genetic disease, the auriculocondylar syndrome (OMIM 602483 and 614669), characterized by severe craniofacial malformations and defects in ear function (Rieder et al., 2012; Gordon et al., 2013; Romanelli Tavares et al., 2015). Studies conducted on Gαi-3 knock-out mice have revealed defects in the immature pre-hearing organ (Beer-Hammer et al., 2018) which depend on the essential role played by Gαi-3 in the control of asymmetric cell division and of the polarity and shaping of the basal cochlea hair cell bundle in the inner ear (Ezan et al., 2013; Mauriac et al., 2017).

The PLCβ3 mutation or depletion also gives rise to several important phenotypes: a homozygous mutation with loss of PLCβ3 function has been linked to spondylometaphyseal dysplasia with corneal dystrophy, an autosomal recessive disease characterized by multiple defects in bone growth and corneal opacity, whose patients’ fibroblasts showed an altered actin cytoskeleton due to increased PIP2 levels (Ben-Salem et al., 2018). In PLCβ3-KO mice, the hyperproliferation of hematopoietic stem cells leads to myeloproliferative neoplasia caused by the lack of suppression of STAT5 signaling mediated by PLCβ3 through the protein phosphatase Shp1 (Xiao et al., 2009).

ηPKC knock-out mice show greater susceptibility to tumor formation in two-stage skin carcinogenesis and show a delayed and defective skin re-epithelialization during cutaneous wound healing, suggesting a role for ηPKC in the maintenance of epithelial tissue architecture (Chida et al., 2003). To what extent these diseases are related to defects in TGN sorting or to other roles of these signaling molecules remains to be clarified.

## Supporting information

Supplementary Information_Di Martino2020

## References

Anitei, M., Stange, C., Czupalla, C., Niehage, C., Schuhmann, K., Sala, P., Czogalla, A., Pursche, T., Coskun, Ü., Shevchenko, A., et al. (2017b). Spatiotemporal Control of Lipid Conversion, Actin-Based Mechanical Forces, and Curvature Sensors during Clathrin/AP-1-Coated Vesicle Biogenesis. Cell Rep 20, 2087–2099.

Baron, C.L., and Malhotra, V. (2002). Role of diacylglycerol in PKD recruitment to the TGN and protein transport to the plasma membrane. Science 295, 325–328.

Ben-Salem, S., Robbins, S.M., Lm Sobreira, N., Lyon, A., Al-Shamsi, A.M., Islam, B.K., Akawi, N.A., John, A., Thachillath, P., Al Hamed, S., et al. (2018). Defect in phosphoinositide signalling through a homozygous variant in PLCB3 causes a new form of spondylometaphyseal dysplasia with corneal dystrophy. J. Med. Genet. 55, 122–130.

Bern, M., Sand, K.M.K., Nilsen, J., Sandlie, I., and Andersen, J.T. (2015). The role of albumin receptors in regulation of albumin homeostasis: Implications for drug delivery. J Control Release 211, 144–162.

Bjarnadóttir, T.K., Schiöth, H.B., and Fredriksson, R. (2005). The phylogenetic relationship of the glutamate and pheromone G-protein-coupled receptors in different vertebrate species. Ann. N. Y. Acad. Sci. 1040, 230–233.

Blüml, K., Schnepp, W., Schröder, S., Beyermann, M., Macias, M., Oschkinat, H., and Lohse, M.J. (1997). A small region in phosducin inhibits G-protein betagamma-subunit function. EMBO J. 16, 4908–4915.

Boncompain, G., Divoux, S., Gareil, N., de Forges, H., Lescure, A., Latreche, L., Mercanti, V., Jollivet, F., Raposo, G., and Perez, F. (2012). Synchronization of secretory protein traffic in populations of cells. Nat. Methods 9, 493–498.

Boutté Y. (2018). Lipids at the crossroad: Shaping biological membranes heterogeneity defines trafficking pathways. PLoS biology, 16, e2005188.

Bräuner-Osborne, H., and Krogsgaard-Larsen, P. (2000). Sequence and expression pattern of a novel human orphan G-protein-coupled receptor, GPRC5B, a family C receptor with a short amino-terminal domain. Genomics 65, 121–128.

Cancino, J., Capalbo, A., Di Campli, A., Giannotta, M., Rizzo, R., Jung, J.E., Di Martino, R., Persico, M., Heinklein, P., Sallese, M., et al. (2014). Control systems of membrane transport at the interface between the endoplasmic reticulum and the Golgi. Dev. Cell 30, 280–294.

Centonze, F. G., Reiterer, V., Nalbach, K., Saito, K., Pawlowski, K., Behrends, C., & Farhan, H. (2019). LTK is an ER-resident receptor tyrosine kinase that regulates secretion. The Journal of cell biology, 218, 2470–2480.

Cheng, Y., and Lotan, R. (1998). Molecular cloning and characterization of a novel retinoic acid-inducible gene that encodes a putative G protein-coupled receptor. J. Biol. Chem. 273, 35008–35015.

Chida, K., Hara, T., Hirai, T., Konishi, C., Nakamura, K., Nakao, K., Aiba, A., Katsuki, M., and Kuroki, T. (2003). Disruption of protein kinase Ceta results in impairment of wound healing and enhancement of tumor formation in mouse skin carcinogenesis. Cancer Res. 63, 2404–2408.

Costa-Mattioli, M., and Walter, P. (2020). The integrated stress response: From mechanism to disease. Science (New York, N.Y.), 368 (6489), eaat5314.

De Matteis, M.A., and Luini, A. (2008). Exiting the Golgi complex. Nat. Rev. Mol. Cell Biol. 9, 273–284.

Di Martino, R., Sticco, L., and Luini, A. (2019). Regulation of cargo export and sorting at the trans-Golgi network. FEBS Lett. 593, 2306–2318.

Díaz Añel, A.M. (2007). Phospholipase C beta3 is a key component in the Gbetagamma/PKCeta/PKD-mediated regulation of trans-Golgi network to plasma membrane transport. Biochem. J. 406, 157–165.

Díaz Añel, A.M., and Malhotra, V. (2005). PKCeta is required for beta1gamma2/beta3gamma2-and PKD-mediated transport to the cell surface and the organization of the Golgi apparatus. J. Cell Biol. 169, 83–91.

Du, F., Nakamura, Y., Tan, T.-L., Lee, P., Lee, R., Yu, B., and Jamora, C. (2010). Expression of snail in epidermal keratinocytes promotes cutaneous inflammation and hyperplasia conducive to tumor formation. Cancer Res. 70, 10080–10089.

Ezan, J., and Montcouquiol, M. (2013). Revisiting planar cell polarity in the inner ear. Semin. Cell Dev. Biol. 24, 499–506.

Farhan H., and Rabouille C. (2011). Signalling to and From the Secretory Pathway. J. Cell Sci. 124, 171–180.

Farhan, H., Kundu, M., and Ferro-Novick, S. (2017). The link between autophagy and secretion: a story of multitasking proteins. Molecular biology of the cell, 28, 1161–1164.

Fogg, V.C., Azpiazu, I., Linder, M.E., Smrcka, A., Scarlata, S., and Gautam, N. (2001). Role of the gamma subunit prenyl moiety in G protein beta gamma complex interaction with phospholipase Cbeta. J. Biol. Chem. 276, 41797–41802.

Garcia-Marcos, M., Ear, J., Farquhar, M.G., and Ghosh, P. (2011). A GDI (AGS3) and a GEF (GIV) regulate autophagy by balancing G protein activity and growth factor signals. Mol. Biol. Cell 22, 673–686.

Giannotta, M., Ruggiero, C., Grossi, M., Cancino, J., Capitani, M., Pulvirenti, T., Consoli, G.M.L., Geraci, C., Fanelli, F., Luini, A., et al. (2012). The KDEL receptor couples to Gαq/11 to activate Src kinases and regulate transport through the Golgi. EMBO J. 31, 2869–2881.

Gissen, P., and Arias, I.M. (2015). Structural and functional hepatocyte polarity and liver disease. J. Hepatol. 63, 1023–1037.

Gordon, C.T., Vuillot, A., Marlin, S., Gerkes, E., Henderson, A., AlKindy, A., Holder-Espinasse, M., Park, S.S., Omarjee, A., Sanchis-Borja, M., et al. (2013). Heterogeneity of mutational mechanisms and modes of inheritance in auriculocondylar syndrome. J. Med. Genet. 50, 174–186.

Guo, Z., Neilson, L.J., Zhong, H., Murray, P.S., Zanivan, S., and Zaidel-Bar, R. (2014). E-cadherin interactome complexity and robustness resolved by quantitative proteomics. Sci Signal 7, rs7.

Hilger, D., Masureel, M., and Kobilka, B. K. (2018). Structure and dynamics of GPCR signaling complexes. Nature structural & molecular biology, 25, 4–12.

Inoue, S., Nambu, T., and Shimomura, T. (2004). The RAIG family member, GPRC5D, is associated with hard-keratinized structures. J. Invest. Dermatol. 122, 565–573.

Jaggi, M., Johansson, S.L., Baker, J.J., Smith, L.M., Galich, A., and Balaji, K.C. (2005). Aberrant expression of E-cadherin and beta-catenin in human prostate cancer. Urol. Oncol. 23, 402–406.

Jamora, C., Yamanouye, N., Van Lint, J., Laudenslager, J., Vandenheede, J.R., Faulkner, D.J., and Malhotra, V. (1999). Gbetagamma-mediated regulation of Golgi organization is through the direct activation of protein kinase D. Cell 98, 59–68.

Jiang, X., Xu, X., Wu, M., Guan, Z., Su, X., Chen, S., Wang, H., and Teng, L. (2018). GPRC5A: An Emerging Biomarker in Human Cancer. Biomed Res Int 2018, 1823726.

Kellokumpu S. (2019). Golgi pH, Ion and Redox Homeostasis: How Much Do They Really Matter?. Frontiers in cell and developmental biology, 7, 93.

Konopka, G., Tekiela, J., Iverson, M., Wells, C., and Duncan, S.A. (2007). Junctional adhesion molecule-A is critical for the formation of pseudocanaliculi and modulates E-cadherin expression in hepatic cells. J. Biol. Chem. 282, 28137–28148.

Kuwano, Y., Adler, M., Zhang, H., Groisman, A., and Ley, K. (2016). Gαi2 and Gαi3 Differentially Regulate Arrest from Flow and Chemotaxis in Mouse Neutrophils. J. Immunol. 196, 3828–3833.

Lambert, A.W., Pattabiraman, D.R., and Weinberg, R.A. (2017). Emerging Biological Principles of Metastasis. Cell 168, 670–691.

Lau, W.W., Chan, A.S., Poon, L.S., Zhu, J., and Wong, Y.H. (2013). Gβγ-mediated activation of protein kinase D exhibits subunit specificity and requires Gβγ-responsive phospholipase Cβ isoforms. Cell Commun. Signal 11, 22.

Liljedahl, M., Maeda, Y., Colanzi, A., Ayala, I., Van Lint, J., and Malhotra, V. (2001). Protein kinase D regulates the fission of cell surface destined transport carriers from the trans-Golgi network. Cell 104, 409–420.

Liu, H., Fang, S., Wang, W., Cheng, Y., Zhang, Y., Liao, H., Yao, H., and Chao, J. (2016). Macrophage-derived MCPIP1 mediates silica-induced pulmonary fibrosis via autophagy. Part Fibre Toxicol 13, 55.

Liu, H., Zhang, Y., Hao, X., Kong, F., Li, X., Yu, J., and Jia, Y. (2016). GPRC5A overexpression predicted advanced biological behaviors and poor prognosis in patients with gastric cancer. Tumour Biol. 37, 503–510.

Liu, S., Ye, D., Wang, T., Guo, W., Song, H., Liao, Y., Xu, D., Zhu, H., Zhang, Z., and Deng, J. (2017). Repression of GPRC5A is associated with activated STAT3, which contributes to tumor progression of head and neck squamous cell carcinoma. Cancer Cell Int. 17, 34.

Luini, A., Mavelli, G., Jung, J., and Cancino, J. (2014). Control systems and coordination protocols of the secretory pathway. F1000prime reports, 6, 88

Luini, A., and Parashuraman, S. (2016). Signaling at the Golgi: sensing and controlling the membrane fluxes. Current opinion in cell biology, 39, 37–42.

MacEwan, D.J. (2002). TNF ligands and receptors--a matter of life and death. Br. J. Pharmacol. 135, 855–875.

Maeda, Y., Beznoussenko, G.V., Van Lint, J., Mironov, A.A., and Malhotra, V. (2001). Recruitment of protein kinase D to the trans-Golgi network via the first cysteine-rich domain. EMBO J. 20, 5982–5990.

Matlin, K.S., and Simons, K. (1983). Reduced temperature prevents transfer of a membrane glycoprotein to the cell surface but does not prevent terminal glycosylation. Cell 34, 233–243.

Matthews, S.A., Rozengurt, E., and Cantrell, D. (1999). Characterization of serine 916 as an in vivo autophosphorylation site for protein kinase D/Protein kinase Cmu. J. Biol. Chem. 274, 26543–26549.

Mauriac, S.A., Hien, Y.E., Bird, J.E., Carvalho, S.D.-S., Peyroutou, R., Lee, S.C., Moreau, M.M., Blanc, J.-M., Geyser, A., Medina, C., et al. (2017). Defective Gpsm2/Gαi3 signalling disrupts stereocilia development and growth cone actin dynamics in Chudley-McCullough syndrome. Nat Commun 8, 14907.

Mellman, I., and Nelson, W. J. (2008). Coordinated protein sorting, targeting and distribution in polarized cells. Nature reviews. Molecular cell biology, 9, 833–845.

Mellman, I., and Warren, G. (2000). The road taken: past and future foundations of membrane traffic. Cell 100, 99–112.

Miller, E.A., Beilharz, T.H., Malkus, P.N., Lee, M.C.S., Hamamoto, S., Orci, L., and Schekman, R. (2003). Multiple cargo binding sites on the COPII subunit Sec24p ensure capture of diverse membrane proteins into transport vesicles. Cell 114, 497–509.

Mironov, A. A., Mironov, A. A., Jr, Beznoussenko, G. V., Trucco, A., Lupetti, P., Smith, J. D., Geerts, W. J., Koster, A. J., Burger, K. N., Martone, M. E., Deerinck, T. J., Ellisman, M. H., & Luini, A. (2003). ER-to-Golgi carriers arise through direct en bloc protrusion and multistage maturation of specialized ER exit domains. Developmental cell, 5, 583–594.

Müsch, A. (2014). The unique polarity phenotype of hepatocytes. Exp. Cell Res. 328, 276–283.

Nieto, M.A., Huang, R.Y.-J., Jackson, R.A., and Thiery, J.P. (2016). EMT: 2016. Cell 166, 21–45.

Pakos-Zebrucka, K., Koryga, I., Mnich, K., Ljujic, M., Samali, A., and Gorman, A. M. (2016). The integrated stress response. EMBO reports, 17, 1374–1395.

Paroutis, P., Touret, N., & Grinstein, S. (2004). The pH of the secretory pathway: measurement, determinants, and regulation. Physiology (Bethesda, Md.), 19, 207–215.

Polishchuk, E.V., Di Pentima, A., Luini, A., and Polishchuk, R.S. (2003). Mechanism of constitutive export from the golgi: bulk flow via the formation, protrusion, and en bloc cleavage of large trans-golgi network tubular domains. Mol. Biol. Cell 14, 4470–4485.

Polishchuk, R.S., Capestrano, M., and Polishchuk, E.V. (2009). Shaping tubular carriers for intracellular membrane transport. FEBS Lett. 583, 3847–3856.

Pulvirenti, T., Giannotta, M., Capestrano, M., Capitani, M., Pisanu, A., Polishchuk, R.S., San Pietro, E., Beznoussenko, G.V., Mironov, A.A., Turacchio, G., et al. (2008). A traffic-activated Golgi-based signalling circuit coordinates the secretory pathway. Nat. Cell Biol. 10, 912–922.

Rieder, M.J., Green, G.E., Park, S.S., Stamper, B.D., Gordon, C.T., Johnson, J.M., Cunniff, C.M., Smith, J.D., Emery, S.B., Lyonnet, S., et al. (2012). A human homeotic transformation resulting from mutations in PLCB4 and GNAI3 causes auriculocondylar syndrome. Am. J. Hum. Genet. 90, 907–914.

Rivera, V.M., Wang, X., Wardwell, S., Courage, N.L., Volchuk, A., Keenan, T., Holt, D.A., Gilman, M., Orci, L., Cerasoli, F., et al. (2000). Regulation of protein secretion through controlled aggregation in the endoplasmic reticulum. Science 287, 826–830.

Robbins, M.J., Charles, K.J., Harrison, D.C., and Pangalos, M.N. (2002). Localisation of the GPRC5B receptor in the rat brain and spinal cord. Mol. Brain Res. 106, 136–144.

Robbins, M.J., Michalovich, D., Hill, J., Calver, A.R., Medhurst, A.D., Gloger, I., Sims, M., Middlemiss, D.N., and Pangalos, M.N. (2000). Molecular cloning and characterization of two novel retinoic acid-inducible orphan G-protein-coupled receptors (GPRC5B and GPRC5C). Genomics 67, 8–18.

Rodriguez-Boulan, E., and Müsch, A. (2005). Protein sorting in the Golgi complex: shifting paradigms. Biochim. Biophys. Acta 1744, 455–464.

Rodriguez-Boulan, E., Kreitzer, G., and Müsch, A. (2005). Organization of vesicular trafficking in epithelia. Nat. Rev. Mol. Cell Biol. 6, 233–247.

Rollins, C.T., Rivera, V.M., Woolfson, D.N., Keenan, T., Hatada, M., Adams, S.E., Andrade, L.J., Yaeger, D., van Schravendijk, M.R., Holt, D.A., et al. (2000). A ligand-reversible dimerization system for controlling protein-protein interactions. Proc. Natl. Acad. Sci. U.S.A. 97, 7096–7101.

Romanelli Tavares, V.L., Gordon, C.T., Zechi-Ceide, R.M., Kokitsu-Nakata, N.M., Voisin, N., Tan, T.Y., Heggie, A.A., Vendramini-Pittoli, S., Propst, E.J., Papsin, B.C., et al. (2015). Novel variants in GNAI3 associated with auriculocondylar syndrome strengthen a common dominant negative effect. Eur. J. Hum. Genet. 23, 481–485.

Sawada, Y., Kikugawa, T., Iio, H., Sakakibara, I., Yoshida, S., Ikedo, A., Yanagihara, Y., Saeki, N., Győrffy, B., Kishida, T., et al. (2020). GPRC5A facilitates cell proliferation through cell cycle regulation and correlates with bone metastasis in prostate cancer. Int. J. Cancer 146, 1369–1382.

Sicart, A., Katan, M., Egea, G., and Sarri, E. (2015). PLCγ1 participates in protein transport and diacylglycerol production triggered by cargo arrival at the Golgi. Traffic 16, 250–266.

Solis, G. P., Bilousov, O., Koval, A., Lüchtenborg, A. M., Lin, C., and Katanaev, V. L. (2017). Golgi-Resident Gαo Promotes Protrusive Membrane Dynamics. Cell, 170, 939–955.e24.

Stoops, E.H., and Caplan, M.J. (2014). Trafficking to the apical and basolateral membranes in polarized epithelial cells. J. Am. Soc. Nephrol. 25, 1375–1386.

Stow, J.L., de Almeida, J.B., Narula, N., Holtzman, E.J., Ercolani, L., and Ausiello, D.A. (1991a). A heterotrimeric G protein, G alpha i-3, on Golgi membranes regulates the secretion of a heparan sulfate proteoglycan in LLC-PK1 epithelial cells. J. Cell Biol. 114, 1113–1124.

Stow, J.L., Sabolic, I., and Brown, D. (1991b). Heterogeneous localization of G protein alpha-subunits in rat kidney. Am. J. Physiol. 261, F831–840.

Subramanian, A., Capalbo, A., Iyengar, N.R., Rizzo, R., di Campli, A., Di Martino, R., Lo Monte, M., Beccari, A.R., Yerudkar, A., Del Vecchio, C., et al. (2019). Auto-regulation of Secretory Flux by Sensing and Responding to the Folded Cargo Protein Load in the Endoplasmic Reticulum. Cell 176, 1461–1476.e23.

Tao, Q., Cheng, Y., Clifford, J., and Lotan, R. (2004). Characterization of the murine orphan G-protein-coupled receptor gene Rai3 and its regulation by retinoic acid. Genomics 83, 270–280.

Tao, Q., Fujimoto, J., Men, T., Ye, X., Deng, J., Lacroix, L., Clifford, J.L., Mao, L., Van Pelt, C.S., Lee, J.J., et al. (2007). Identification of the retinoic acid-inducible Gprc5a as a new lung tumor suppressor gene. J. Natl. Cancer Inst. 99, 1668–1682.

Tapia, D., Jiménez, T., Zamora, C., Espinoza, J., Rizzo, R., González-Cárdenas, A., Fuentes, D., Hernández, S., Cavieres, V.A., Soza, A., et al. (2019). KDEL receptor regulates secretion by lysosome relocation-and autophagy-dependent modulation of lipid-droplet turnover. Nat Commun 10, 735.

Thompson, B.D., Jin, Y., Wu, K.H., Colvin, R.A., Luster, A.D., Birnbaumer, L., and Wu, M.X. (2007). Inhibition of G alpha i2 activation by G alpha i3 in CXCR3-mediated signaling. J. Biol. Chem. 282, 9547–9555.

Van IJzendoorn, S.C.D., Van Der Wouden, J.M., Liebisch, G., Schmitz, G., and Hoekstra, D. (2004). Polarized membrane traffic and cell polarity development is dependent on dihydroceramide synthase-regulated sphinganine turnover. Mol. Biol. Cell 15, 4115–4124.

Walter, P., and Ron, D. (2011). The unfolded protein response: from stress pathway to homeostatic regulation. Science 334, 1081–1086.

Wendeler, M.W., Paccaud, J.-P., and Hauri, H.-P. (2007). Role of Sec24 isoforms in selective export of membrane proteins from the endoplasmic reticulum. EMBO Rep. 8, 258–264.

Wilson, B.S., Komuro, M., and Farquhar, M.G. (1994). Cellular variations in heterotrimeric G protein localization and expression in rat pituitary. Endocrinology 134, 233–244.

Wilson, P.D. (2011). Apico-basal polarity in polycystic kidney disease epithelia. Biochim. Biophys. Acta 1812, 1239–1248.

Xiao, W., Hong, H., Kawakami, Y., Kato, Y., Wu, D., Yasudo, H., Kimura, A., Kubagawa, H., Bertoli, L.F., Davis, R.S., et al. (2009). Tumor suppression by phospholipase C-beta3 via SHP-1-mediated dephosphorylation of Stat5. Cancer Cell 16, 161–171.

Xu, J., Tian, J., and Shapiro, S.D. (2005). Normal lung development in RAIG1-deficient mice despite unique lung epithelium-specific expression. Am. J. Respir. Cell Mol. Biol. 32, 381–387.

Ye, X., Tao, Q., Wang, Y., Cheng, Y., and Lotan, R. (2009). Mechanisms underlying the induction of the putative human tumor suppressor GPRC5A by retinoic acid. Cancer Biol. Ther. 8, 951–962.

Yeaman, C., Ayala, M.I., Wright, J.R., Bard, F., Bossard, C., Ang, A., Maeda, Y., Seufferlein, T., Mellman, I., Nelson, W.J., et al. (2004). Protein kinase D regulates basolateral membrane protein exit from trans-Golgi network. Nat. Cell Biol. 6, 106–112.

Young, A., Jiang, M., Wang, Y., Ahmedli, N.B., Ramirez, J., Reese, B.E., Birnbaumer, L., and Farber, D.B. (2011). Specific interaction of Gαi3 with the Oa1 G-protein coupled receptor controls the size and density of melanosomes in retinal pigment epithelium. PLoS ONE 6, e24376.

Zanetti, G., Pahuja, K.B., Studer, S., Shim, S., and Schekman, R. (2011). COPII and the regulation of protein sorting in mammals. Nat. Cell Biol. 14, 20–28.

Zhou, H., and Rigoutsos, I. (2014). The emerging roles of GPRC5A in diseases. Oncoscience 1, 765–776.

