## Supplementary Information_Di Martino2020 for "Autoregulatory circuit regulating basolateral cargo export from the TGN: role of the orphan receptor GPRC5A in PKD signaling and cell polarity": SUPPLEMENTARY INFORMATION_DI MARTINO ET AL. 2020.pdf

Contents:

1. SUPPLEMENTARY FIGURE S1- S4

2. MATERIALS AND METHODS

3. TABLES T1-T3

#### Supplementary Figure 1

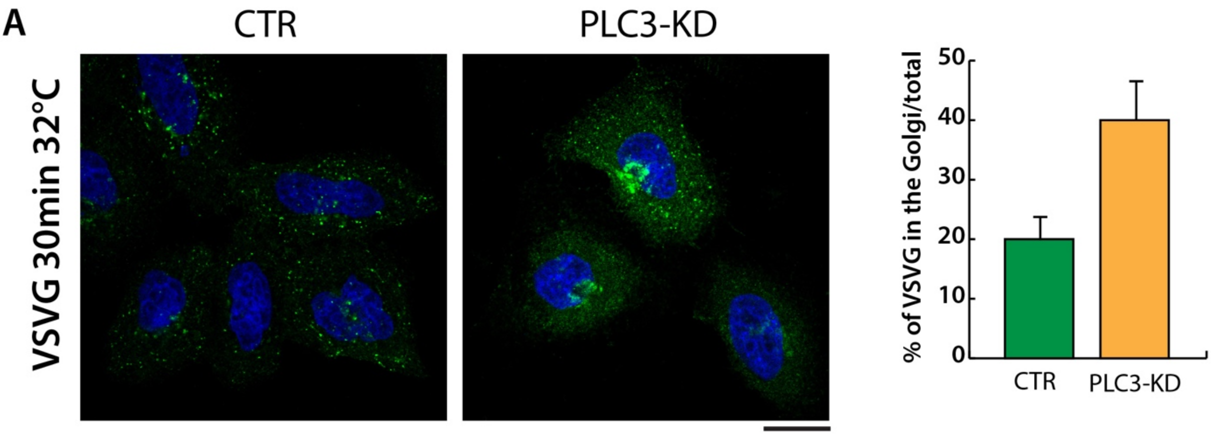


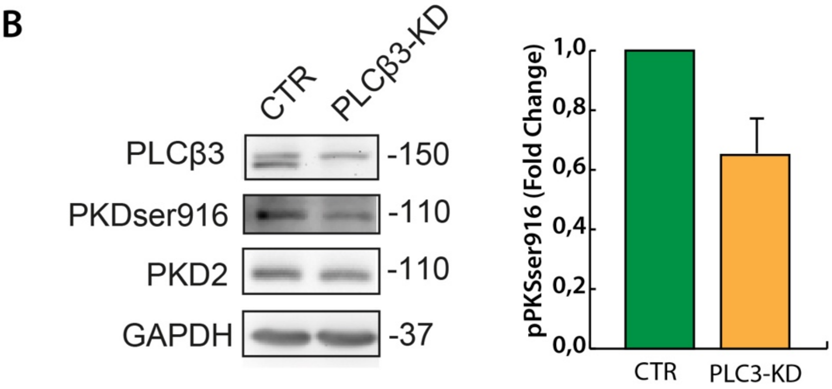


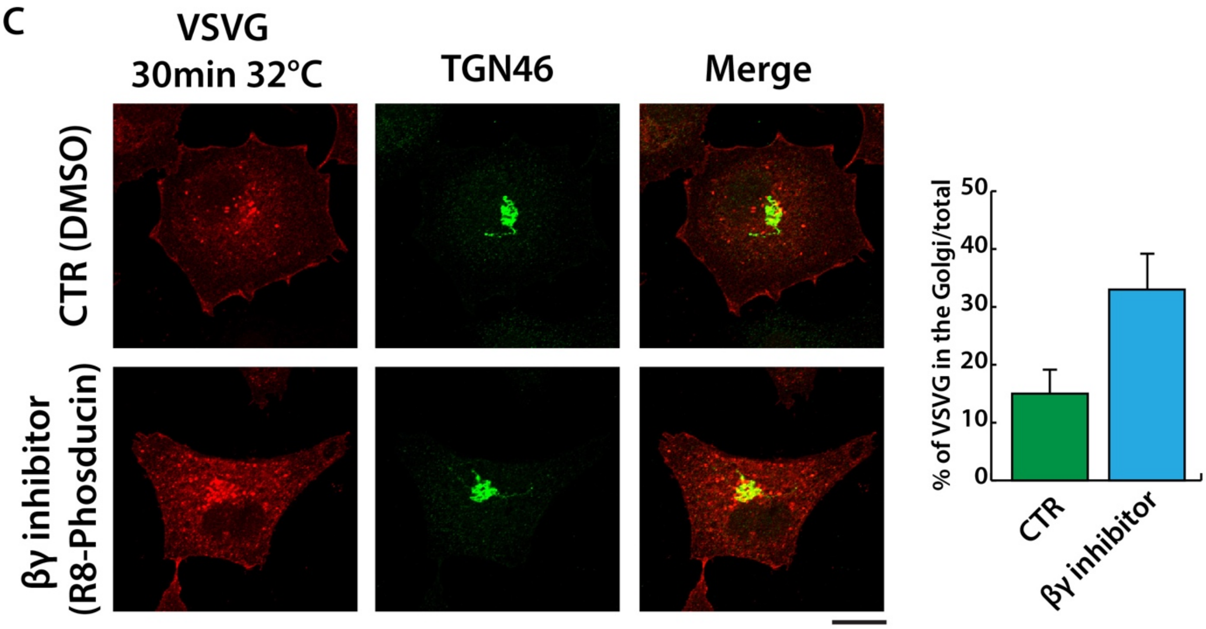

**Supplementary Figure 1: PLCβ3 depletion and Gβγ inhibition block VSVG export from the TGN to the PM. A)** IF analysis of VSVG localization in PLCβ3-KD and control HeLa cells at the time point of 30 minutes after the release of the 20°C block. **B)** WB analysis of pPKDser916 levels in steady state PLCβ3-KD and control HeLa cells. **C)** IF analysis of VSVG localization in HeLa cells treated with Gβγ-inhibitor (R8-phosducin 50μM) or vehicle (DMSO) at the time point of 60 minutes after the release of the 20°C block.

### Supplementary Figure 2

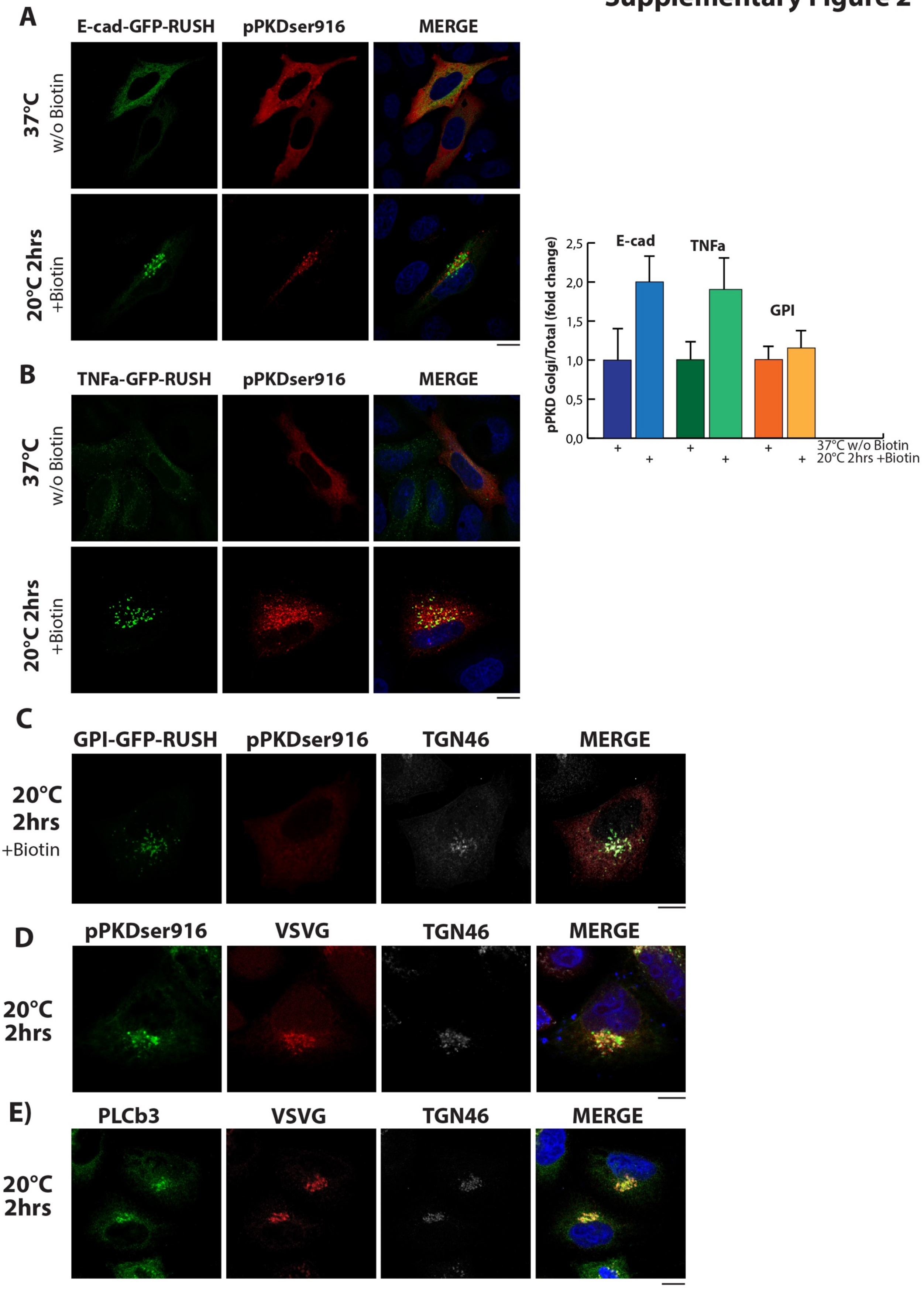

**Supplementary Figure 2: Basolateral but not apical cargoes accumulation in the TGN induce pPKDser916 activation on TGN membranes in HeLa cells. A-B-C)** IF analysis of overexpressed PKD1-GST labeled with pPKDser916 antibody during the accumulation in the TGN (20°C 2hrs) of E-cadherin-RUSH-GFP (basolateral), TNFa-RUSH-GFP (basolateral) and GPI-GFP-RUSH (apical) respectively. **D-E)** IF Co-localization analysis of pPKDser916 (overexpressed PKD1-GST) and endogenous PLCb3 respectively with both VSVG and TGN46 during TGN accumulation at 20°C for 2 hours (Scale bars: 10nm).

#### Supplementary Figure 3:

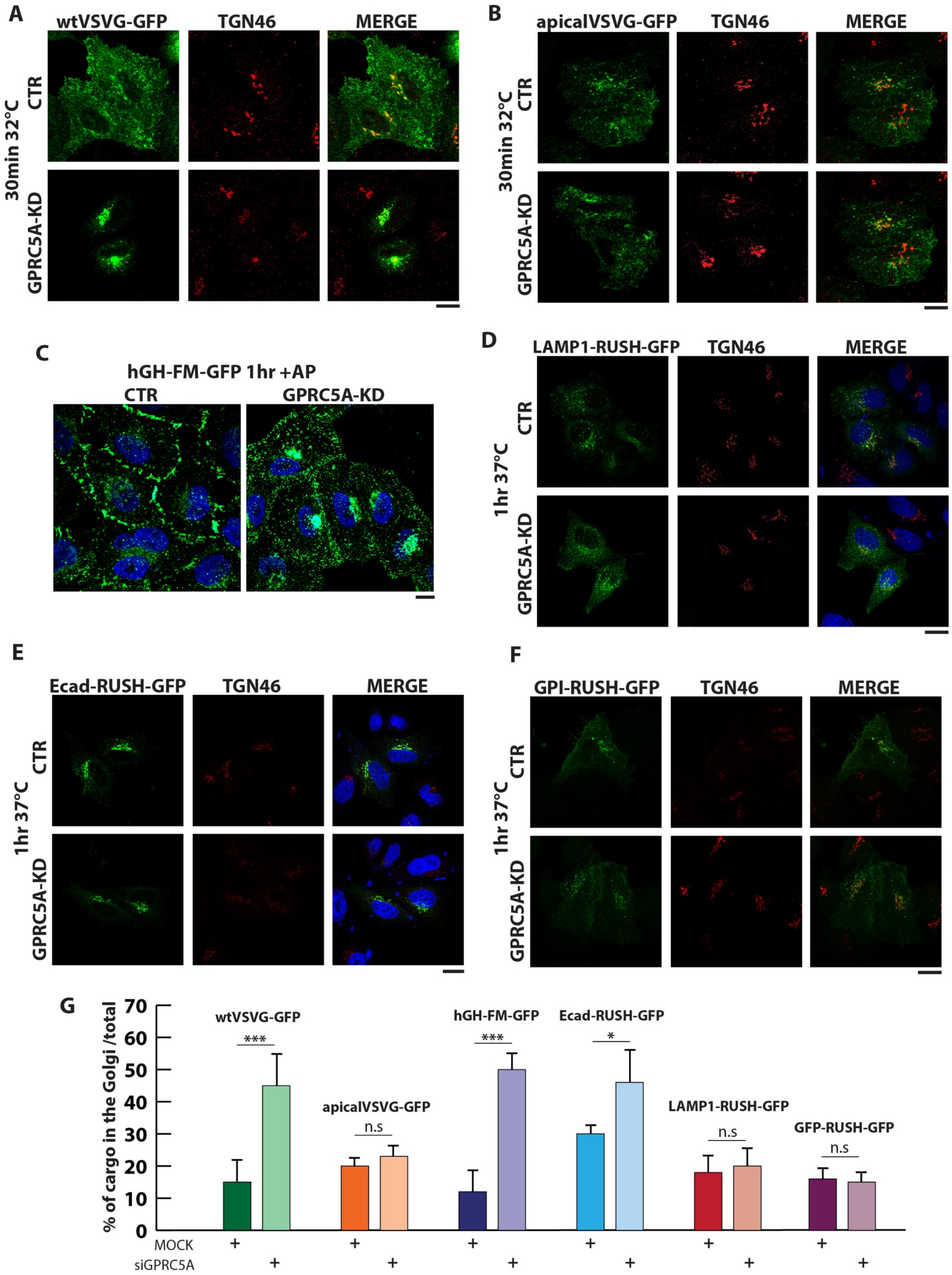

**Supplementary Figure 3: GPRC5A depletion affects basolateral cargo arrival to the plasma membrane, but not apical or lysosomal cargoes. A-B)** IF analysis of the localization of GFP constructs for VSVG wild type (basolateral) and apical mutant respectively at the time point of 30 minutes at 32°C after the release of the 20°C block, in HeLa control and GPRC5A-KD cells. **C)** IF analysis of the stably transfected human Growth Hormone (hGH) –FM-GFP in control and GPRC5A-KD HeLa cells at the time point of 1 hour at 37°C after the addition of the AP drug to depolymerize the FM domains. **D-E-F)** IF analysis of the localization of GFP-RUSH constructs for LAMP1 (lysosomal), E-cadherin (basolateral) and GPI (apical) at the time point of 1 hour at 37°C after the release of the 20°C block in control and GPRC5A-KD HeLa cells (scale bars: 10nm). **G**) Quantification of the percentage of the above-mentioned cargoes in the Golgi compared to the total.

#### Supplementary Figure 4

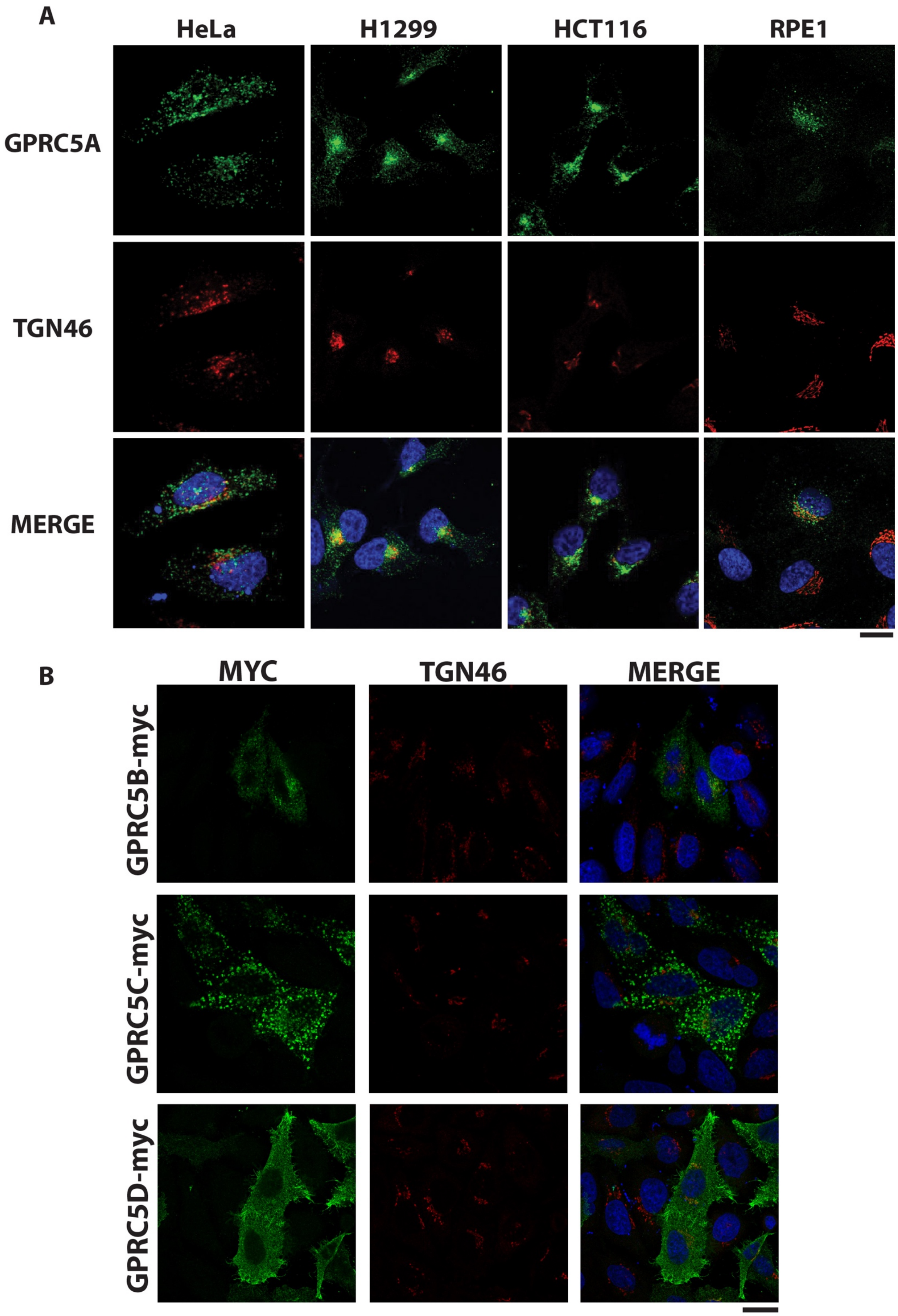

**Supplementary Figure 4**: **Immunofluorescence localization of endogenous GPRC5A in different human cell lines and of MYC-tagged GPRC5-B-C-D in HeLa cells. A)** IF staining for endogenous GPRC5A has been performed on steady state HeLa (cervical cancer), H1299 (non-small cell lung carcinoma), HCT116 (colorectal carcinoma) and RPE1 (hTERT-immortalized retinal pigmented epithelial cell line). The TGN marker TGN46 colocalizes with GPRC5A. **B)** IF localization of overexpressed GPRC5B-myc, GPRC5C-myc and GPRC5D-myc proteins in steady state HeLa cells.

2.

#### Materials and Methods

##### Cell lines

HeLa-M (human cervical cancer cells, female origin) were obtained from the ATCC and grown in RPMI-1640 supplemented with 10% FCS. HeLa-M cells stably transfected with hGH-FM-GFP (female origin) were a kind gift from A Peden (Gordon et al., 2010) and grown in DMEM supplemented with 10% FCS. HepG2 (human hepatocyte carcinoma cells) were obtained from the ATCC and grown in DMEM supplemented with 10% FCS. For experiments with the RUSH constructs, HeLa-M cells were grown in DMEM supplemented with 10% FCS without biotin. No mycoplasma contamination was observed in cell lines. The cell lines have not been authenticated.

##### Transfection

HeLa-M cells were transfected with plasmid vectors using the TransIT-LT1 or Lipofectamine LTX reagents. Only cells expressing moderate levels of the respective plasmids were chosen for IF analyses. All siRNA treatments were carried out using a pool of siRNAs and Oligofectamine. Expression or knockdown efficiencies (> 85%) were checked after every experiment either by indirect immunofluorescence or immunoblotting. Each individual siRNA in the pool was also tested for knockdown and in transport assays with similar results. A list of siRNA sequences used in this study can be found in Table S2.

##### Cargo transport pulses

Six synchronizable secretory cargos were used in this study – one temperature sensitive (ts) cargo ts045-VSVG (VSVG), and five different molecular trap cargoes (SBP-GFP-E-Cadherin, SBP-GFP-TNFalpha, SBP-GFP-GPI, SBP-GFP-LAMP1, and hGH-FM-GFP). VSVG viral infection was performed at 32°C for 1 h in FCS-free DMEM-HEPES medium. Cells then were washed 3 times with PBS and kept for 3 h at 40°C in DMEM-HEPES supplemented with 10% FCS to accumulate unfolded VSVG in the ER. For ts045-VSVG-GFP transfections, cells were incubated with the plasmid for 16 h at 40°C in an incubator with 5%CO2. An ER to Golgi/TGN transport pulse was induced by shifting cells to 32°C for 20 min or later time points as indicated (Mironov et al., 2003, Nishimura and Balch, 1997), while a TGN to plasma membrane pulse was induced by incubating the cells at 20°C for 2hrs and then shifting the temperature to 32°C for the indicated time points. Molecular trap cargo RUSH constructs were transfected into HeLa cells cultured in

DMEM supplemented with 10% FCS without biotin at 37°C. A traffic pulse was induced by treating cells with biotin (40 μg/mL) for the indicated times (Boncompain et al., 2012). Similarly, synchronous hGH-FM-GFP release from the ER of HeLa cells stably transfected with the cargo was performed by adding DD solubilizer (2μM) to the cells for the times indicated in each figure (Gordon et al., 2010). Cycloheximide (CHX; 50 μg/mL) was added 30 min prior to all traffic pulse experiments to ensure that a majority of the secretory cargo could be monitored and the response it generates in the cell was specific.

**Targeted phosphoproteomics by antibody microarray and analysis**

HeLa cells infected with the VSVG virus were subjected to an ER block (40°C 3 h, see above) or a traffic pulse (32°C 20min). Cells were lysed in lysis buffer (50 mM HEPES, pH 7.4, 100 mM NaCl, 0.5% NP-40, 30 mM NaF, 2 mM Na3O4V, 60 mM β-glycerophosphate, 5 mM EDTA, 5 mM EGTA, and the protease inhibitor cocktail from Roche) and the protein concentrations were quantified using BCA kits (Pierce). Protein samples were frozen at −80°C before being subjected to an antibody microarray (KAM-880 array) and data analysis, which was performed by the Kinexus service (System proteomic company, Vancouver, Canada). The array monitors changes in the expression levels and phosphorylation states of signaling proteins which includes 518 pan-specific antibodies (for protein expression) and 359 phospho-site-specific antibodies (for changes in phosphorylation). The resultant changes are expressed as percentages of change with respect to the control (CFC) and as Z-factors. Changes ≥ 50% CFC and Z ratio ≥ +1.00 (hyper phosphorylation) or ≤ −1.00 (dephosphorylation) were considered as real significant changes. Only proteins whose phosphorylation status differed after the folding pulse were considered, analyzed by bioinformatic tools such as Ingenuity Pathway Analysis, DAVID-KEGG pathway, STRING and PhosphoSite plus and built into signaling pathway maps or networks manually using Adobe Illustrator (Adobe systems).

**Inhibitor and peptide treatments**

All inhibitors and cell permeant peptides were tested for purity and efficacy before use. In most experiments, cells were pretreated with them for 1 h in serum-free media unless otherwise specified. Gö6976 (Calbiochem) treatment: HeLa cells were treated with 10uM Go6976 (PKD Inhibitor) at 20°C from 1h before temperature shift at 32°C until the end of the traffic pulse in culture medium without FCS and buffered by 20 mM HEPES, pH 7.4. R8-Phosducin (aa 215-232) peptide treatment (G beta/gamma inhibitor): R8-Phosducin has been synthesized in collaboration with Dr. Petra Hienkel from University of Basel, according to Blum et al, 1997.

HeLa cells were treated with 50 μM R8Phos(aa215-232) at 20°C, 1h minutes before the temperature shift until the end of the traffic pulse experiment as previously described for PKD inhibitor.

**Generation of GPRC5A siRNA resistant construct**

GPRC5A siRNA resistant construct was generated by site-directed mutagenesis. The oligonucleotide primers used for the mutagenesis reaction were:

5’-AGAACAGAGCCTACAGCCAGGAAGAAATCACTCAAGGTTTTGA-3’
5’-ACCTTGAGTGATTTCTTCCTGGCTGTAGGCTCTGTTCTCCAC-3’.

PCR conditions for mutagenesis reactions were as follows: a single denaturation step (95°C, 5 min) was performed and followed by 20 cycles of denaturation (95°C, 1 min), annealing (60°C, 1 min) and elongation (68°C, 30 min). Then a final elongation step of 10 min at 72°C was performed. The PCR reaction was carried out in a 50 µl final volume including the following components: 50 ng of DNA template, 500 µM of each dNTPs, 1.5x PfuTurbo Cx Hotstart DNA Polymerase buffer, 0.2 µM of each primer and 5 U of PfuTurbo Cx Hotstart DNA Polymerase. Following completion of mutagenesis reactions, parental plasmid DNA was digested by DpnI treatment (20 U/mutagenesis reaction) for 1 h at 37°C. Then DNA was precipitated, resuspended in sterile water and transformed into chemical competent E. coli TOP-10 cells.

**Confocal microscopy**

Images were acquired using a Zeiss LSM710 using a 40x or 63 × oil-immersion objective (1.4 NA) with identical settings for each channel throughout single experiments. Indirect immunofluorescence: indirect immunofluorescence (IF) was performed as follows: cells grown on coverslips were washed in phosphate-buffered saline (PBS) and fixed in freshly prepared PBS supplemented with 4% paraformaldehyde (Electron Microscopy Sciences, Hatfield, USA) for 10 min at room temperature (RT) for most experiments. Cells were permeabilized and blocked in blocking buffer (0.05% saponin, 0.5% BSA, ammonium chloride 50mM, in PBS1X) for 30 min at room temperature. Primary antibodies were incubated for 1h at RT or overnight at 4°C in blocking buffer. Cells were subsequently labeled with appropriate Alexa 488/568/647-tagged fluorescent-conjugated secondary antibodies (Invitrogen). Samples on glass coverslips were mounted on glass microscope slides (Carlo Erba, Italy) using Mowiol (20 mg Mowiol dissolved in 80 ml PBS, stirred O/N and centrifuged for 30 min at 12,000x g).

Image processing: for figure presentation only, the images were channel-separated, with each channel shown as color or grayscale image after correcting for contrast using Image-J (NIH) or Adobe Photoshop CS3 (Adobe Systems). In some cases, the contrast was inverted, and the levels adjusted to facilitate the observation of dim structures.

**SDS–PAGE and Immunoblotting**

The cells were washed three times with ice-cold PBS and collected immediately at 4°C in phospho-RIPA lysis buffer (1% Triton X-100, 20mM MOPS pH 7.0, 30 mM NaF, 1 mM $Na_3O_4V$, 60 mM β-glicerophosphate, 5 mM EDTA, 2 mM EGTA and Proteases cocktail inhibitor from Roche. Cell lysates were centrifuged at 14,000 rpm for 15 min at 4°C to the eliminate detergent insoluble pellet. The supernatant was immediately processed for SDS–PAGE and immunoblotting with antibodies.

**Immunoprecipitation**

Total lysates were prepared using IP lysis buffer (150 mM NaCl, 25mM Tris-HCl pH 7.5, 1% Triton-X, 10mM Na3VO4, 40mM β-glycerophosphate, 10mM NaF and protease cocktail inhibitor from Roche). The protein concentrations were quantified and 1mg of protein was used for immunoprecipitation with antibodies conjugated to either Protein A Sepharose or magnetic dynabeads. For GiGTP IP only the immunoprecipitation protocol has been performed accordingto the manufacture protocol (G alpha i Activation Assay Kit -80301- Product Manual).

**TCA/ACETONE precipitation**

Cell media was replaced with serum-free media before starting the experiment, then 1ml of media was collected for each condition and centrifuged at 14000 RPM at 4 °C for 10 minutes to eliminate cell debris. TCA was added to the supernatant to a final concentration of 20–25% and the proteins were allowed to precipitate on ice for at least 2 hours to overnight. Subsequently, proteins were centrifuged for 30 minutes at 14000 RPM at 4 °C. The pellet obtained was washed with ice-cold acetone and centrifuged again for 15 minutes at 14000 RPM at 4°C. This step was repeated three times. Residual acetone was removed by air drying, and at the end the pellet was dissolved in sample buffer with β-mercaptoethanol and subjected to SDS-PAGE for analysis.

#### Membrane/cytosol fractionation

Briefly, HeLa cells were washed with PBS and detached using trypsin, and then centrifuged for 5 minutes at 1500 RPM. The cell pellet is re-suspended in 0.1X PBS with a protease inhibitor cocktail and incubated on ice for 10 minutes. The cells were then manually disrupted using the pestle of a Dounce Homogenizer, allowing the cells to break but leaving nuclei intact. 10X PBS was added in order to have 1XPBS concentration in the final volume and then centrifuged at 4°C for 20 min at 2500 rpm to pellet the nuclei. The supernatant containing cytosol and membranes was ultra-centrifugated at 100.000g for 1 hour at 4°C. The supernatant obtained from this spin contains the cytosol fraction, while the pellet contains the membranes.

#### Quantitative fluorescence image analysis

Quantitative analysis was performed using the Image J software. In brief, to calculate the amount of cargo in the TGN after a traffic pulse, the integrated intensity fluorescence was measured for each cell area and for the TGN area of that cell and the TGN/Total ratio was calculated.

#### Densitometric Analysis

Western blots were acquired using the Chemidoc Imaging System (Bio-Rad) and the exposure times were varied to obtain appropriate signal intensities of protein bands. Each of the bands were quantitated using the Image-J gel analysis tool.

#### Statistical Analysis

All the statistical analysis was performed using Graphpad Prism software. P Value corresponds to (*** 0.005, ** 0.001, * 0.05) and ns corresponds to not significant. Error bars correspond to standard deviation (S.D). Students unpaired t test was used throughout the manuscript unless otherwise stated.

**TABLE T1: List of primary and secondary antibodies used for WB, IF and IP**

| Name | Catalog Number | Source |
|---|---|---|
| Anti-pPKDser916 | 2051 | Cell Signaling |
| Anti-pERK | 9101 | Cell Signaling |
| Anti-pSTAT1 | 9167 | Cell Signaling |
| Anti-ZO1 | 5406 | Cell Signaling |
| Anti-Myc Tag | 2272 | Cell Signaling |
| Anti-GAPDH | SC-32233 | Santa Cruz |
| Anti-PKD2 | SC-100415 | Santa Cruz |
| Anti-Gi3 | SC-262 | Santa Cruz |
| Anti-ERK | SC-94 | Santa Cruz |
| Anti-Calnexin | SC-23954 | Santa Cruz |
| Anti-VSVG | A190-131A | Bethil |
| Anti-GPRC5a | HPA007928 | Sigma |
| Anti-pCAMK4 | SAB4504122 | Sigma |
| Anti VSVG (P5D4) | SAB4200695 | Sigma |
| Anti-active GiGTP | 26901 | New East Bioscience |
| Anti E-Cadherin | 610182 | BD Bioscience |
| Anti-GFP | ab137827 | Abcam |
| Anti-human TGN46 | AHP500G | BioRad/AbD-Serotec |
| Anti-PLCb3 | GTX111100 | Genetex |
| Anti-GST rabbit polyclonal | N/A | kindly provided by Antonella De Matteis's Lab |
| Anti-human Alpha 1 anti trypsin | A0012 | DAKO |
| Anti-human albumin | A0001 | DAKO |
| Alexa fluor secondary antibodies (for IF) | / | Thermo Fisher Scientific |
| HRP conjugated secondary antibodies (for WB) | / | Calbiochem |

**TABLE T2: List of siRNAs used in this study**

| Gene name | Sequence |
|---|---|
| **GPRC5A** | 5' GCCUACUCUCAAGAGGAAA 3'<br>5' GCAAUGGCCUGAAAUCCAA 3'<br>5' CAAAGACUAUGAAGUAAAG 3'<br>5' GCUCAUGCUUCCUGACUUU 3' |
| **G alpha i 3** | 5' CCAAGGAGAUCUAUACUCA 3'<br>5' GAUGAUCGACCGCAACUUA 3'<br>5' GGGAAUAUCAGCUCAAUGA 3'<br>5' GAAUAUCCCAGUCUAACUA 3' |
| **PLCb3** | 5' GGAGUAAGUUCAUCAAAUG 3'<br>5' GAACAAAUGCUUCGAGAUG 3'<br>5' CCAUGGAGUUUGUGGAAUA 3'<br>5' GAGGAGGAACCCUUCAUUA 3' |
| **GPRC5C** | 5' CAGGUGAUGGGCAGUGCCA 3'<br>5' GUAGAGGUCAUCAUCAAUA 3'<br>5' GGAUCGUCAUGUAUACUUA 3'<br>5' UCUGGCGGCUCACGUCUUU 3' |

**TABLE T3: List of plasmids used in this study**

| Plasmid name | Source |
|---|---|
| GPRC5A-Myc | Origene technologies Catalog No. RC200118 |
| siRNAresistant-GPRC5A-Myc | This study (see Methods fro details) |
| GPRC5B-Myc | Origene technologies Catalog No. RC205201 |
| GPRC5C-Myc | Origene technologies Catalog No. RC211081 |
| GPRC5D-Myc | Origene technologies Catalog No. RC210138 |
| VSVG-RFP | Kindly provided by Dr. Jorge Cancino |
| E-cadherin(RUSH)-GFP | Kindly provided by Dr. Frank Perez |
| GPI(RUSH)-GFP | Kindly provided by Dr. Frank Perez |
| TNFalpha(RUSH)-GFP | Kindly provided by Dr. Frank Perez |
| Lamp1(RUSH)-GFP | Kindly provided by Dr. Frank Perez |
| PKD-GST | Kindly provided by Dr. Vivek Malhotra |
| C1a-GST | Kindly provided by Dr. Vivek Malhotra |
